# Cap-independent co-expression of dsRNA-sensing and NF-κB pathway inhibitors enables tunable self-amplifying RNA expression with reduced immunotoxicity

**DOI:** 10.1101/2024.09.24.614636

**Authors:** Tony K.Y Lim, Anne Ritoux, Luke W. Paine, Larissa Ferguson, Tawab Abdul, Laura J. Grundy, Ewan St. John Smith

## Abstract

Self-amplifying RNA (saRNA) has the potential to provide durable, non-integrating transgene expression for transient gene therapy. However, its auto-replicative nature mimics viral infection, triggering innate immune responses that shut down cap-dependent translation, degrade cellular mRNA, induce cell death, and release cytokines. In non-immunotherapy applications, this immune activation is undesirable as it limits transgene expression, induces unintended changes in host gene expression, depletes transfected cells, and promotes inflammation—ultimately undermining therapeutic outcomes. Moreover, the use of exogenous immune suppressants to mitigate these effects often increases treatment complexity and the risk of unintended systemic side effects. To address these challenges, we developed a strategy to encode broad-spectrum innate immune suppression directly within saRNA. This approach leverages cap-independent translation to bypass saRNA-triggered translation shutdown, enabling the expression of multiple inhibitors targeting diverse double-stranded RNA-sensing and inflammatory signalling pathways. In mouse primary fibroblast-like synoviocytes—key cells in joint pathologies—this strategy eliminates the need for external immune inhibitors, reduces cytotoxicity and antiviral cytokine secretion, and enables sustained transgene expression that can be controlled with a small-molecule antiviral. Together, these findings support the development of reversible ‘immune-evasive saRNA’ constructs for transient gene therapy applications that avoid persistent immune activation and eliminate the need for external immune suppressants.

**GRAPHICAL ABSTRACT:** 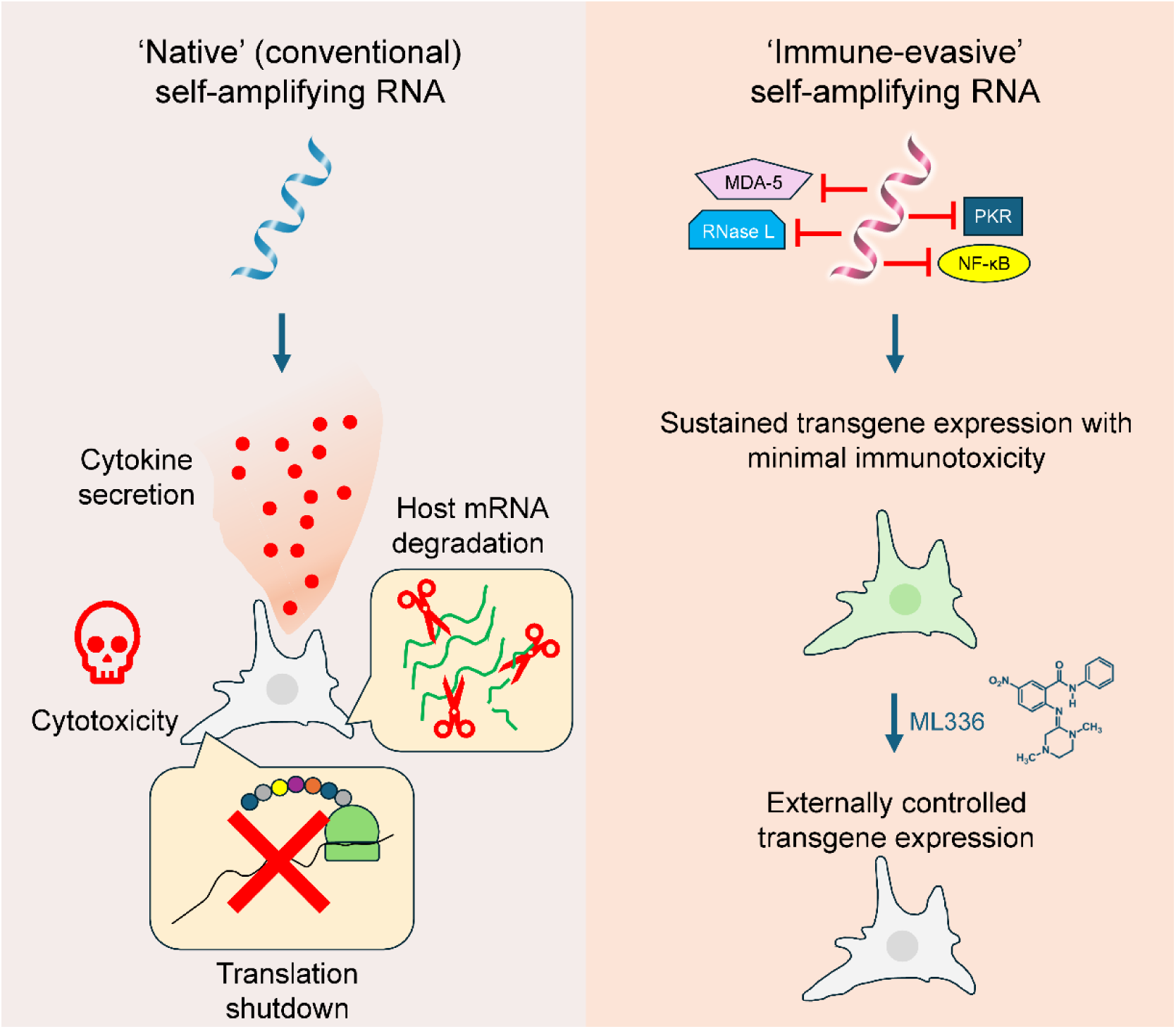

## INTRODUCTION

Self-amplifying RNA (saRNA) is a promising platform for transient gene therapy due to its ability to encode large therapeutic payloads and achieve sustained, non-integrating protein expression.^1^ Upon cellular internalization, saRNA utilizes host machinery to synthesize an RNA-dependent RNA polymerase (RdRp), which drives replication of the saRNA genome.^2^ This process involves the production of a negative-strand RNA intermediate that serves as a template for both genomic and subgenomic RNA synthesis.^3^ Subgenomic RNA, encoding the gene of interest, is transcribed in excess of the genomic RNA, leading to robust and sustained protein expression.^4^

However, the self-replication of saRNA also generates double-stranded RNA (dsRNA) intermediates, formed both during the synthesis of negative-strand RNA and during the transcription of positive-strand RNA from the negative-strand template.^3^ These dsRNA intermediates are potent activators of cytosolic pattern recognition receptors, triggering innate immune responses.^5,6^ This recognition results in shutdown of cap-dependent translation,^7–10^ degradation of cellular mRNA,^6,11^ induction of programmed cell death,^6,10,12–16^ and release of cytokines and chemokines characteristic of viral infection.^9,17,18^ In non-immunotherapy applications such as transient gene therapy, these responses are particularly problematic, as they limit transgene expression, reshape the host transcriptional landscape in ways that may undermine therapeutic effects, deplete transfected cells, and induce inflammation.

Strategies to mitigate saRNA-mediated innate immune responses have included the incorporation of modified nucleotides^19^, removal of dsRNA contaminants,^9,20^ and co-delivery of non-replicating mRNA encoding viral innate immune inhibiting proteins.^21–23^ While these approaches can reduce initial innate immune responses to saRNA transfection, they fall short in addressing the continuous immune activation triggered by dsRNA intermediates that arise during saRNA replication.^2^ Sequence evolution of the RdRp shows some promise in reducing these responses, but even engineered RdRps still induce significant innate immune activation.^6,24^ Encoding viral immune inhibitors within the saRNA construct using 2A self-cleaving peptides can improve protein expression *in vitro* but has limited impact on cytokine responses, despite reductions in NF-κB and IRF3 activation.^18^ Although these strategies temper saRNA-induced innate immune responses to levels appropriate for immunotherapy applications—where some immune activation is beneficial^25^—they fall short of the stringent requirements for non-immunotherapy contexts, where therapeutic outcomes can be compromised by immune activation. In such cases, exogenous immunosuppressants can mitigate saRNA-induced immune responses and enable effective transgene expression,^9,22,26,27^ but this approach complicates treatment regimens and increases the risk of unintended side effects from systemic immune suppression.

To address these challenges, we developed a fully saRNA-based approach that mitigates innate immune responses triggered by saRNA replication, incorporating several key innovations. First, innate immune inhibitory proteins are expressed via cap-independent translation, bypassing the cap-dependent translation shutdown commonly triggered by saRNA. Cap-dependent translation requires the presence of a 5′ cap structure on mRNA, which recruits translation initiation factors and ribosomes.^28^ By contrast, cap-independent translation uses internal ribosome entry sites (IRES) to recruit ribosomes directly to the mRNA, enabling translation even when cap-dependent pathways are inhibited.^29^ Second, by simultaneously targeting multiple dsRNA-sensing and inflammatory signalling pathways, our method provides a more comprehensive suppression of innate immune responses than single target approaches. Third, encoding innate immune inhibitors directly within the saRNA ensures continuous protection against persistent innate immune activation caused by dsRNA intermediates during saRNA replication, eliminating the need for exogenous immunosuppressants.

We evaluated this approach in mouse primary fibroblast-like synoviocytes (FLS), a key cell type in the pathology of joint diseases.^30,31^ To facilitate this evaluation, we developed a microplate assay for straightforward, longitudinal monitoring of saRNA translation control and cytotoxicity in live cells. Our results demonstrate that this strategy effectively reduces saRNA-induced cytotoxicity and cytokine production while enabling sustained cap-dependent transgene expression, which can be reversed with a small-molecule antiviral. These results pave the way for ‘immune-evasive’ saRNA constructs that enable durable and externally controllable transgene expression with minimal innate immune activation, all without the need for exogenous immunosuppressants.

## RESULTS

### Development of a microplate assay for longitudinal monitoring of translational control and cell number

Since saRNA can induce cap-dependent translation shutdown and cell death, we developed a microplate assay to simultaneously monitor cap-independent translation, cap-dependent translation, and cell number over time. In this assay, primary mouse FLS (Supplemental Figure S1) were plated in 6-well plates and transfected with saRNA constructs that used the Venezuelan equine encephalitis virus (VEEV) RdRp to self-replicate. These constructs featured a dual-fluorescence design, with EGFP expressed through encephalomyocarditis virus (EMCV) IRES-mediated, cap-independent translation, and mScarlet3 expressed through subgenomic cap-dependent translation. This dual-reporter system enabled assessment of the effects of saRNA on translational control (Figure 1a). Spectral overlap between EGFP and mScarlet3 was corrected using linear unmixing (Supplemental Figure S2a–c).

**Figure 1.**
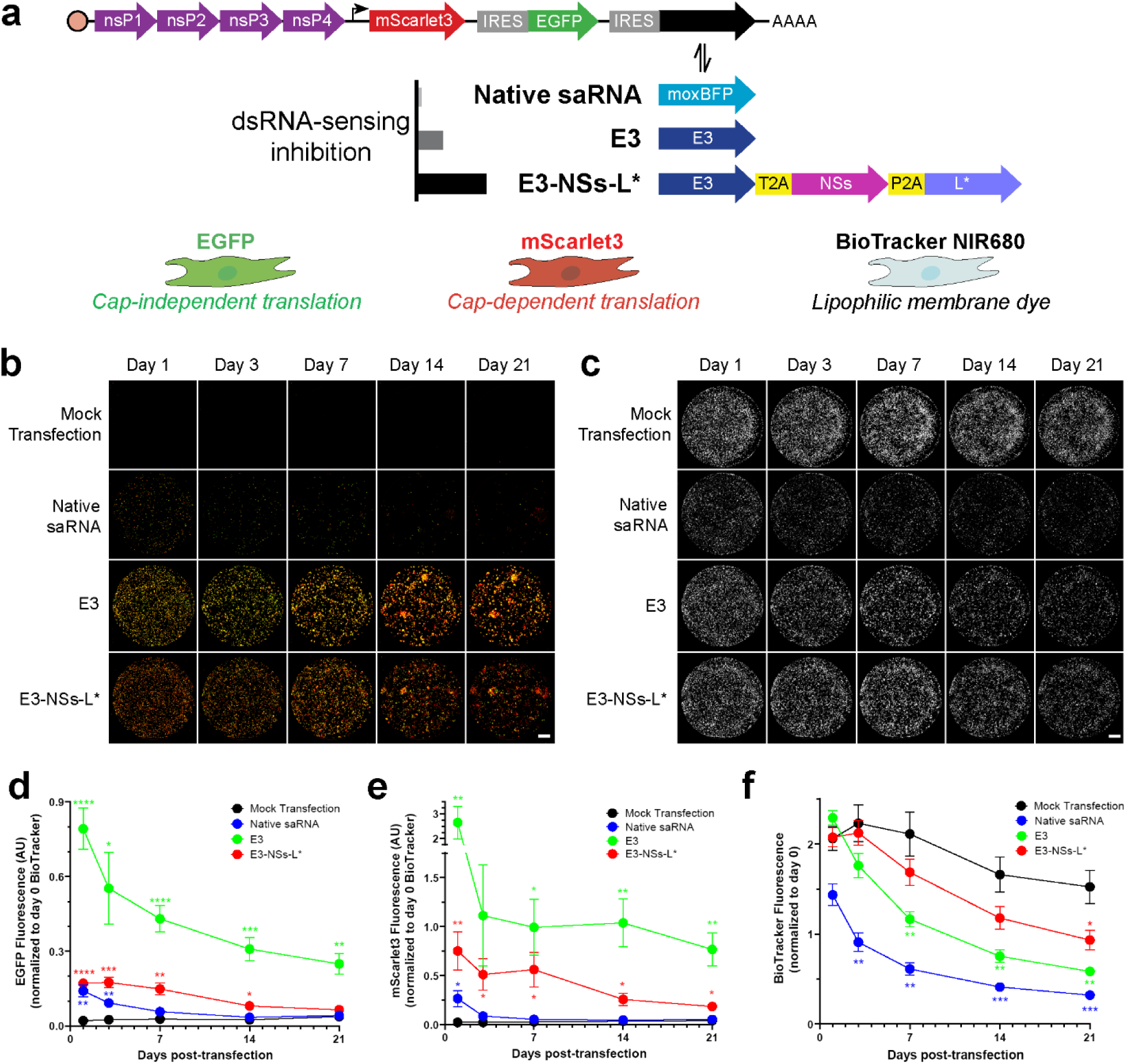
Differential effects of moderate and strong dsRNA-sensing pathway inhibition on saRNA transgene expression and cell loss **a,** Schematic of the native saRNA, E3, and E3-NSs-L* constructs designed to inhibit dsRNA-sensing pathways and report saRNA transgene expression. The native saRNA construct lacks dsRNA-sensing inhibitors. The E3 construct expresses vaccinia virus E3, a pleiotropic inhibitor of dsRNA sensing, expected to provide moderate inhibition. The E3-NSs-L* construct expresses vaccinia virus E3, and additionally includes Toscana virus NSs and Theiler’s virus L*, which target the PKR and OAS/RNase L pathways, expected to provide strong inhibition. EGFP is expressed via an IRES to report cap-independent translation, while a subgenomic promoter (depicted with an angled arrow) enables transcription of an RNA transcript that expresses mScarlet3 via cap-dependent translation. saRNA constructs were transfected into primary mouse FLS, which were labeled with BioTracker to monitor cell number. **b,** Representative images of EGFP (green) and mScarlet3 (red) expression in FLS transfected with native saRNA, E3, or E3-NSs-L* over 3 weeks. Scale bar = 5 mm. **c,** Representative images of FLS transfected with the same constructs, showing BioTracker intensity over time. Scale bar = 5 mm. **d,** Quantification of EGFP fluorescence over time (n = 11 biological replicates). The E3 construct provided the greatest EGFP expression, while the E3-NSs-L* construct showed intermediate levels. Statistical significance of treatment effects at each time point compared to mock transfection was determined by two-way RM ANOVA with Greenhouse–Geisser correction and Dunnett’s multiple comparisons test. **e,** Quantification of mScarlet3 fluorescence over time (n = 11 biological replicates). The E3 construct provided the greatest mScarlet3 expression, while the E3-NSs-L* showed intermediate levels. Statistical significance of treatment effects at each time point compared to mock transfection was determined by two-way RM ANOVA with Greenhouse–Geisser correction and Dunnett’s multiple comparisons test. **f,** Quantification of BioTracker fluorescence over time (n = 11 biological replicates). The native saRNA and E3 constructs reduced BioTracker fluorescence, indicating cell loss. Statistical significance of treatment effects at each time point compared to mock transfection was determined by two-way RM ANOVA with Greenhouse–Geisser correction and Dunnett’s multiple comparisons test. For all statistical reporting, *P < 0.05, **P < 0.01, ***P < 0.001 and ****P < 0.0001. Data are presented as mean ± SEM. For panels (d-f): Data were normalized to starting cell number, indicated by BioTracker intensity on day 0, prior to transfection. The mock transfection control data is also presented in Figure 5c-e. Acronyms: saRNA, self-amplifying RNA; dsRNA, double-stranded RNA; nsP, Venezuelan equine encephalitis virus non-structural protein; IRES, encephalomyocarditis virus internal ribosome entry site; moxBFP, monomeric oxidizing environment-optimized blue fluorescent protein; E3, Vaccinia virus E3 protein; NSs, Toscana virus non-structural NSs protein; L*, Theiler’s murine encephalomyelitis virus L* protein; T2A, Thosea asigna virus 2A peptide; P2A, porcine teschovirus-1 2A peptide; PKR, protein kinase R; OAS, oligoadenylate synthase; AUC, area under the curve.

To monitor cell number, FLS were stained with BioTracker NIR680, a lipophilic carbocyanine membrane dye that enables long-term fluorescent labeling of cells.^32^ Carbocyanine dyes like BioTracker are weakly fluorescent in aqueous solution but exhibit high fluorescence when integrated into lipid bilayers.

Consequently, mock transfection with Lipofectamine MessengerMAX—a lipid transfection agent— enhanced BioTracker fluorescence compared to non-transfected cells, likely due to an interaction with the lipid components of Lipofectamine (Supplemental Figure S3a and b). Despite this interaction, BioTracker remains effective for indicating cell number, as treatment of mock-transfected cells with the apoptosis inducer staurosporine^33^ reduced BioTracker fluorescence. The BioTracker results were consistent with those obtained using the viability dye calcein AM (Supplemental Figure S3c and d), with the added advantages of eliminating the need for additional staining and wash steps and avoiding spectral overlap with red and green fluorescent proteins. This assay allows for seamless longitudinal monitoring of cap-independent translation, cap-dependent translation, and cell number using microplate readers or imagers.

### Design of saRNA constructs for inhibiting multiple dsRNA-sensing pathways

To attenuate saRNA-induced innate immune responses, we first started by designing constructs that inhibit dsRNA-sensing pathways. Cells detect cytosolic dsRNA through several pathways, including RIG-I,^34^ melanoma differentiation-associated protein 5 (MDA-5),^35^ protein kinase R (PKR),^36^ and oligoadenylate synthase (OAS)/RNase L pathways.^37^ To broadly inhibit these pathways, we designed an saRNA construct, referred to as ‘E3’, that expresses the vaccinia virus E3 protein (Fig. 1a). E3 protein is a pleiotropic inhibitor that binds and sequesters dsRNA, effectively blocking multiple dsRNA-sensing pathways.^38^

Given the key roles of PKR in cap-dependent translation shutdown and OAS/RNase L in cellular mRNA degradation in response to dsRNA, we sought to further inhibit these pathways in a second construct. This construct includes Toscana virus NSs, a ubiquitin ligase that promotes PKR degradation,^39^ and Theiler’s virus L*, which inhibits RNase L.^40^ To enable co-expression of these three proteins—E3, NSs, and L*—we incorporated 2A “self-cleaving” peptide sequences between the proteins, allowing the polyprotein to be cleaved into separate proteins during translation. As prior experiments showed that using multiple identical 2A peptides can reduce protein expression,^41^ we selected non-identical 2A peptides for this construct. This construct, named ‘E3-NSs-L*’, is expected to provide a more comprehensive inhibition of dsRNA-sensing pathways than the E3 construct.

Because translation shutdown is one of the major cellular responses to saRNA replication, we expressed these dsRNA-sensing pathway inhibitors using ECMV IRES-mediated, cap-independent translation.

During translation shutdown, global cap-dependent mRNA translation initiation is limited, freeing ribosomes that would otherwise be used for translating capped mRNAs,^42^ thereby enhancing cap-independent translation.^43^ By leveraging this mechanism, IRES-mediated expression of dsRNA-sensing pathway inhibitors ensures their levels increase as translation shutdown intensifies. In contrast, encoding these inhibitors via cap-dependent translation is expected to be less effective, as translation shutdown would lead to reduced inhibitor expression.

As a control, we designed a ‘native saRNA’ construct that expresses a blue fluorescent protein (moxBFP) as a visual marker of IRES functionality in place of inhibitors of dsRNA-sensing pathways. Importantly, moxBFP expression did not overlap spectrally with the EGFP or mScarlet3 channels (Supplemental Figure S2b).

### Moderate dsRNA-sensing pathway inhibition enables high transgene expression at the cost of cell loss, while strong inhibition preserves cell number at lower expression levels

Following transfection of saRNA constructs, we monitored BioTracker, EGFP, and mScarlet3 fluorescence over 3 weeks (Figure 1b and c). Inhibiting dsRNA-sensing pathways with viral proteins significantly enhanced saRNA transgene expression (Figure 1d and e). Interestingly, the E3 construct produced the highest levels of EGFP and mScarlet3 expression, surpassing both native saRNA and the E3-NSs-L* construct. Native saRNA yielded low levels of transgene expression, while E3-NSs-L* showed intermediate expression.

Achieving durable transgene expression depends equally on preserving cell viability and maximizing transgene expression, as constructs that induce cell death undermine the goal of sustained transgene expression. Transfection of FLS with native saRNA caused both immediate and long-term reductions in cell number, as indicated by BioTracker intensity (Figure 1f). While the E3 construct initially maintained cell number, BioTracker signals gradually diminished over time. In contrast, the E3-NSs-L* construct provided sustained protection against the decline in cell number. Area under the curve (AUC) analysis quantified these cumulative effects, confirming that E3-NSs-L* significantly preserved cell numbers compared to E3 (Figure 1—figure supplement 1a). CellTag-based normalization assays on day 2 post-transfection further corroborated these findings, revealing a stepwise increase in protection against cell loss with greater dsRNA-sensing pathway inhibition across the three constructs (Figure 1—figure supplement 1b). Together, these results indicate that while the E3 construct enhances saRNA transgene expression, it is associated with increased cell loss, highlighting the advantage of E3-NSs-L* for applications requiring sustained, non-cytotoxic gene expression.

### Inhibiting saRNA replication mitigates saRNA-induced cytotoxicity

*In vitro* transcription of saRNA often generates dsRNA byproducts, which can activate innate immune pathways and contribute to cytotoxicity.^20,44^ Consequently, it remains unclear whether saRNA-induced cell loss arises primarily from these byproducts or from cytosolic dsRNA generated during saRNA replication.

To investigate this, native saRNA was transfected into FLS in the presence of 10 μM ML336, a potent inhibitor of the VEEV RdRp that mediates self-amplification.^45^ Cultures that did not receive ML336 exhibited reduced calcein AM viability staining the day after transfection. In contrast, ML336 treatment significantly rescued cell viability (Figure 1—figure supplement 1c), suggesting that saRNA-induced cell loss results primarily from replication-associated dsRNA, rather than transcriptional byproducts.

### Inhibiting dsRNA-sensing pathways reduces saRNA-induced cytotoxicity and improves cell viability

saRNA is known to induce programmed cell death,^6,10,12–16^ which often leads to cell detachment.^46^ Thus reductions in BioTracker signal following saRNA treatment are likely indicative of this process. However, since BioTracker is a lipophilic membrane dye, the reduction in signal may not necessarily correlate with cell viability or cytotoxicity. In addition to long-term cell tracking, lipophilic membrane dyes like BioTracker are often used to stain extracellular vesicles, which bud off from the plasma membrane.^47^ During cellular stress, increased production of extracellular vesicles^48^ could deplete the membrane-associated BioTracker dye from cells, resulting in a lower detectable signal. Therefore, the observed reduction in BioTracker signal could reflect enhanced extracellular vesicle production rather than cell death.

To provide more definitive evidence that inhibiting dsRNA-sensing pathways protects against saRNA-induced cytotoxicity, we conducted a longitudinal assay with annexin V staining over 6 days (Figure 2a), followed by calcein AM staining on day 7 (Figure 2b). Annexin V, a membrane-impermeable protein, binds to phosphatidylserine, which translocates to the extracellular side of the cell membrane during early apoptosis, enabling the detection of apoptotic cells with intact membranes.^49^ If the membrane is compromised, annexin V can enter cells and stain intracellular phosphatidylserine, indicating late apoptosis or necrosis.^50^ Calcein AM, a viability dye, accumulates in metabolically active cells with intact membranes, serving both as a marker of live cell number and an indicator of membrane integrity.^51^

**Figure 2.**
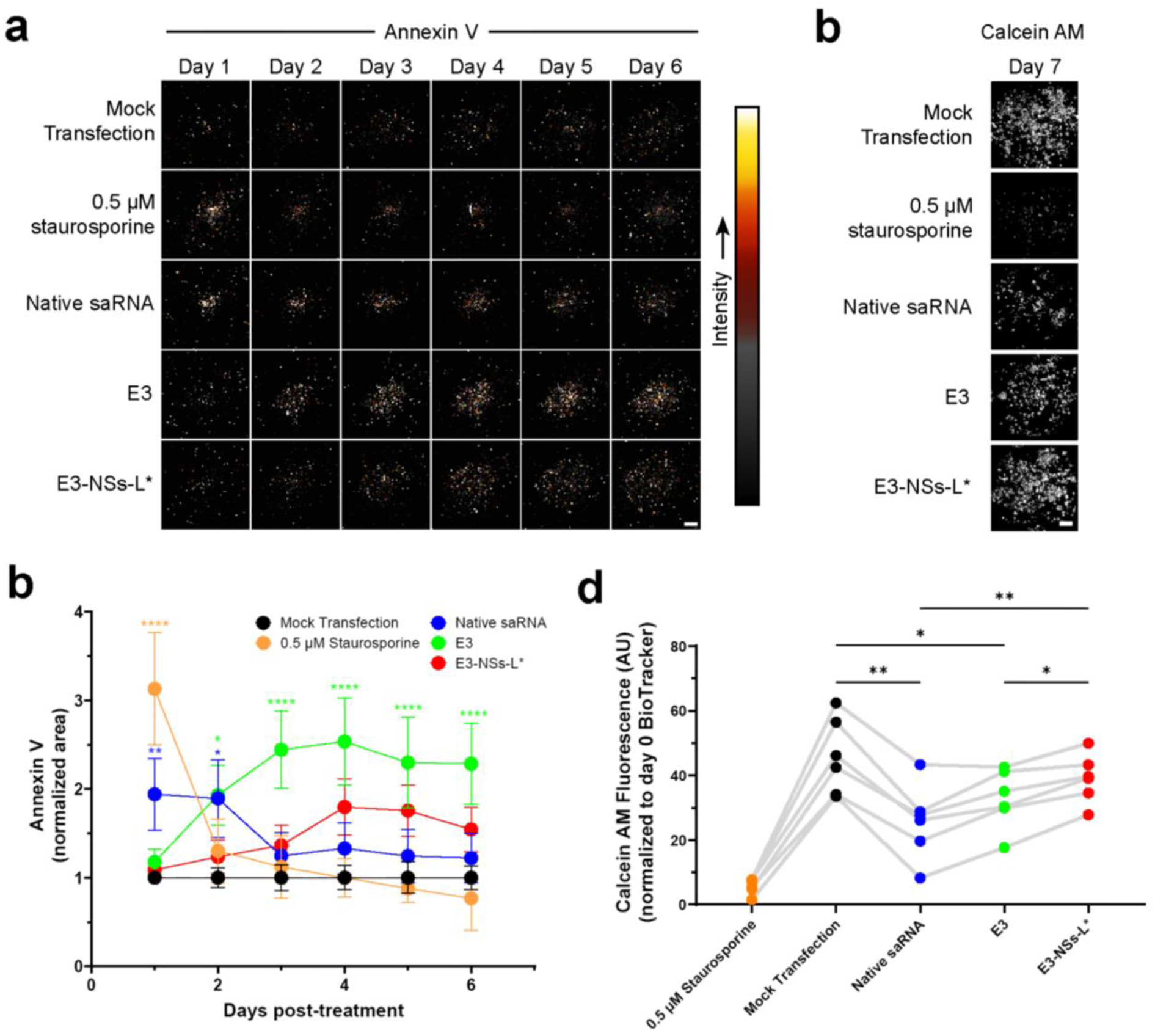
saRNA induces increased phosphatidylserine staining and reduced viability, which is prevented by E3-NSs-L* **a**, Representative cropped images of Annexin V-CF800 staining, indicating phosphatidylserine exposure or loss of membrane integrity, performed daily over 6 days using a microplate imager. Scale bar = 1.5 mm. **b**, Representative cropped images of calcein AM staining, indicating viability, on day 7 post-treatment. Scale bar = 1.5 mm. **c**, Annexin V staining, quantified as the area of positive pixels determined using Li thresholding (n = 6 biological replicates). Staurosporine, native saRNA, and the E3 construct significantly increased annexin V staining, while E3-NSs-L* did not. Data are normalized to the average of the mock transfection group. Statistical significance of treatment effects at each time point compared to mock transfection was determined using two-way RM ANOVA with Bonferroni’s multiple comparisons test. Data are presented as mean ± SEM. **d**, Calcein AM intensity measured on day 7 post-treatment (n = 6 biological replicates). Native saRNA, and E3 significantly reduced cell viability compared to mock transfection, while E3-NSs-L* did not. Cell viability in the E3-NSs-L* group was significantly higher than the E3 group. Connecting lines indicate responses from the same biological replicate. All groups differed significantly from staurosporine; these comparisons are omitted from the figure for clarity due to the large number of statistical comparisons. Data are normalized to cell number on day 0 as determined by BioTracker staining before transfection. Statistical significance was determined by one-way RM ANOVA with Greenhouse–Geisser correction and Tukey’s multiple comparisons test comparing all groups. saRNA constructs used are shown in Fig. 1a. For all statistical reporting, *P < 0.05, **P < 0.01, ***P < 0.001 and ****P < 0.0001.

The apoptosis inducing agent staurosporine caused a transient increase in annexin V staining that decreased after the first day (Figure 2c), along with a marked reduction in calcein AM fluorescence compared to mock transfection (Figure 2d), observations that are consistent with extracellular phosphatidylserine translocation and/or increased membrane permeability. Similarly, native saRNA induced a temporary increase in annexin V staining during the first two days post-transfection (Figure 2c) accompanied by reduced calcein AM staining (Figure 2d). Co-expression of E3 prevented the initial increase in annexin V staining, but elevated it from day 2 onward, and also resulted in significantly reduced calcein AM intensity (Figure 2c and d). In contrast, E3-NSs-L* had no significant effect on annexin V or calcein AM staining compared to mock transfection, supporting the idea that broader inhibition of dsRNA-sensing pathways, achieved by combining E3, NSs and L*, is more effective than E3 alone in preventing saRNA-induced cell death and preserving cell viability.

### dsRNA-sensing pathway inhibition prevents eIF2α phosphorylation, but fails to affect saRNA-induced reductions in eIF4E phosphorylation levels

After establishing how moderate and strong inhibition of dsRNA-sensing pathways affects saRNA-induced cytotoxicity, we next investigated how these pathways modulate translational control at the molecular level. Eukaryotic initiation factor 2 alpha (eIF2α) is a central regulator of translation initiation; its phosphorylation serves as a cellular stress response to limit cellular cap-dependent protein synthesis.^42^ Using in-cell western assays, we found that native saRNA transfection increased eIF2α phosphorylation, but both E3 and E3-NSs-L* effectively blocked this phosphorylation (Figure 3a), consistent with previous reports of vaccinia virus E3’s inhibitory effect on eIF2α phosphorylation.^21,23^ Interestingly, E3-NSs-L* also increased total eIF2α levels, an effect not observed with the E3 construct (Figure 3b).

**Figure 3.**
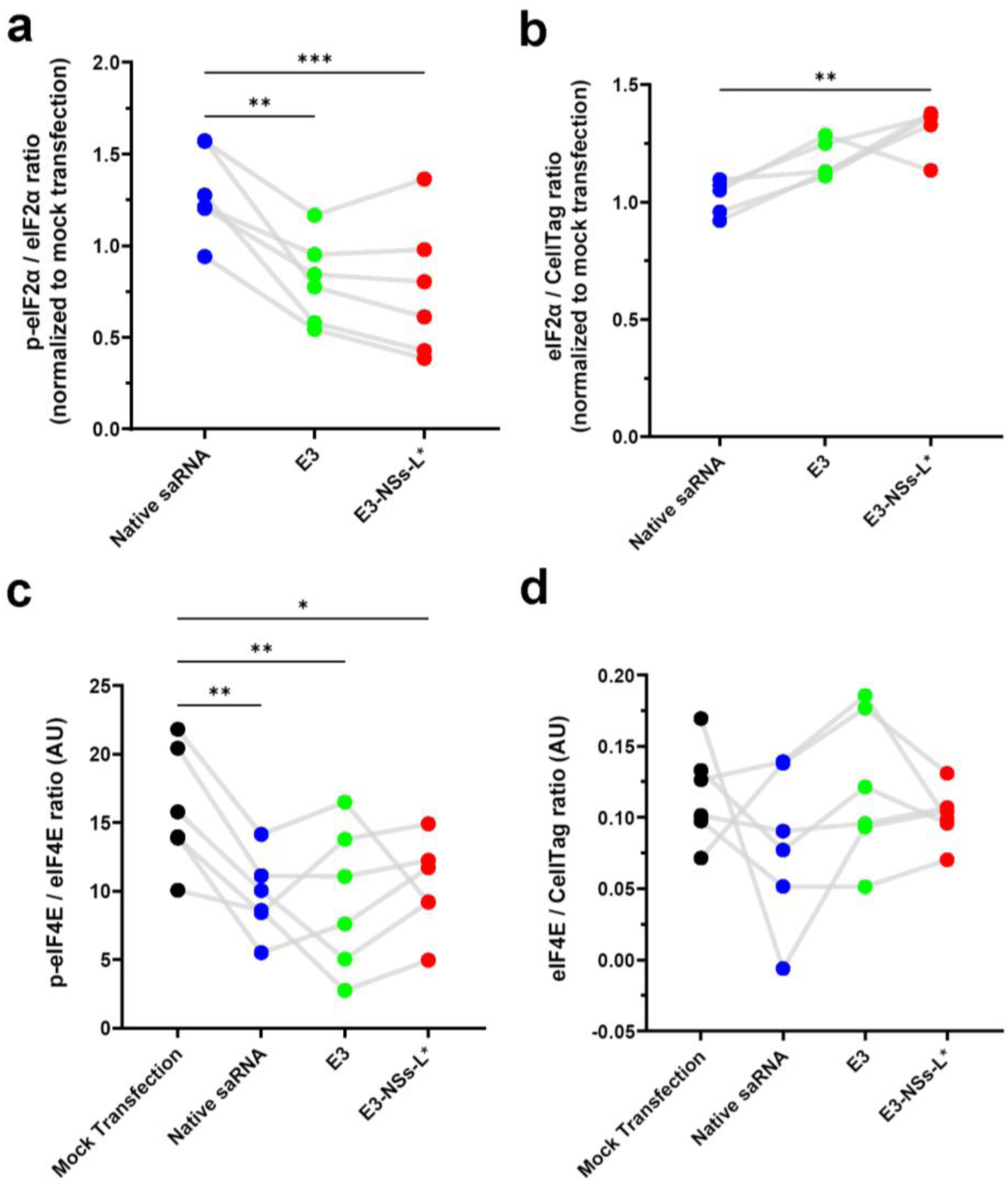
dsRNA-sensing pathway inhibitors suppress saRNA-induced eIF2α phosphorylation but do not affect saRNA-induced reductions in eIF4E phosphorylation **a,** Phosphorylated eIF2α levels examined day 2 post-transfection by in-cell western assay (n = 6 biological replicates). Both E3 and E3-NSs-L* constructs significantly reduced eIF2α phosphorylation. Data are presented as fold-change relative to mock-transfected cells. Statistical significance was determined by one-way RM ANOVA with Tukey’s multiple comparisons test. **b,** eIF2α levels examined day 2 post-transfection by in-cell western assay (n = 5 biological replicates). E3-NSs-L* significantly increased total eIF2α levels compared to native saRNA transfection. Data are presented as fold-change relative to mock-transfected cells. Statistical significance was determined by one-way RM ANOVA with Tukey’s multiple comparisons test. **c,** Phosphorylated eIF4E levels examined day 2 post-transfection by in-cell western assay (n = 6 biological replicates). All saRNA constructs tested significantly reduced eIF4E phosphorylation levels compared to mock transfection. Statistical significance was determined by one-way RM ANOVA with Tukey’s multiple comparisons test. **d,** eIF4E levels examined day 2 post-transfection by in-cell western assay (n = 6 biological replicates). One-way RM ANOVA revealed no significant differences between groups (F(3,15) = 1.207, P = 0.3410). saRNA constructs used are shown in Figure 1a. Connecting lines indicate responses from the same biological replicate. Statistical significance is indicated as follows: *P < 0.05, **P < 0.01, ***P < 0.001. The mock transfection control data are also used in Figure 6a–d.

We next examined the impact of saRNA transfection on eIF4E, another key regulator of translation initiation.^52^ Unlike eIF2α, whose phosphorylation was selectively inhibited by the E3 and E3-NSs-L* constructs, all constructs—including those that inhibit dsRNA-sensing pathways—reduced eIF4E phosphorylation to a similar extent (Figure 3c), while total eIF4E levels remained unchanged (Figure 3d). These results indicate that dsRNA-sensing pathway inhibition does not prevent the saRNA-induced reduction in eIF4E phosphorylation, suggesting that cap-dependent translation remains broadly suppressed across all conditions.

Given that PKR phosphorylates eIF2α upon activation by dsRNA,^7^ we next assessed PKR levels following saRNA transfection. Consistent with previous reports,^6^ native saRNA transfection led to increased PKR levels (Figure 3—figure supplement 1a). The E3 construct produced a similar increase, whereas co-expression of E3-NSs-L* mitigated this effect. This finding aligns with the established role of the Toscana virus NSs protein as a ubiquitin ligase that targets PKR for degradation.^39^

### Host mRNA degradation contributes to enhanced transgene expression under moderate dsRNA-sensing pathway inhibition

The increased transgene expression observed with the E3 construct compared to E3-NSs-L* cannot be attributed to differences in eIF2α or eIF4E-mediated translational control. Phosphorylation levels of both factors were comparable between constructs (Figure 3a and c), indicating that the enhanced transgene expression associated with the E3 construct arises from alternative mechanisms.

Upon detection of dsRNA, OAS enzymes synthesize 2′-5′-oligoadenylates, which activate RNase L, a ribonuclease that degrades single-stranded RNA. This leads to widespread host mRNA cleavage,^53^ impaired nuclear export of transcripts,^54^ and global arrest of protein synthesis.^55^ Several RNA viruses, including dengue and Zika, exploit this process by localizing their transcripts within replication organelles, shielding them from RNase L activity while maintaining efficient translation.^54,56^ The VEEV-derived saRNA used in this study also replicates within these organelles,^57^ likely protecting its transcripts from RNase L– mediated degradation in a similar fashion.

Consequently, RNase L activation by the E3 construct may preferentially deplete host mRNAs, thereby reducing competition for the ribosomal machinery and enhancing the relative translation of protected saRNA transcripts. In contrast, E3-NSs-L* expresses Theiler’s virus L*, an RNase L inhibitor,^40^ preventing host mRNA degradation and forcing saRNA to compete with cellular mRNA for ribosome access.

To evaluate whether RNase L activation contributes to the greater transgene expression observed with the E3 construct compared to E3-NSs-L*, we assessed ribosomal RNA (rRNA) integrity as a surrogate measure of RNase L activity.^58^ Total RNA was extracted from FLS transfected with either E3 or E3-NSs-L*, and evaluated using the RNA integrity number (RIN) algorithm, an automated Bayesian learning method that quantifies rRNA integrity.^59^ Cells transfected with E3 exhibited significantly lower RIN values compared to those transfected with E3-NSs-L* (Figure 3—figure supplement 1b). As prior studies have shown that RNase L activation primarily depletes host mRNA levels before extensive rRNA degradation,^53,55,60^ these findings support the hypothesis that RNase L activation and subsequent reductions in cellular mRNA occur with the E3 construct but not with E3-NSs-L*.

We further evaluated global protein synthesis using surface sensing of translation (SUnSET),^61,62^ in which FLS were pulsed with puromycin, and newly synthesized peptides were detected via anti-puromycin immunoreactivity (Figure 3—figure supplement 1c). Transfection with native saRNA led to a marked reduction in protein synthesis, consistent with translation shutdown. Co-expression of E3 partially restored translation; however, synthesis rates remained significantly lower than in mock transfected cells, likely due to continued RNase L–mediated depletion of host mRNAs despite relief from translation shutdown. In contrast, transfection with E3-NSs-L* resulted in protein synthesis rates comparable to those of mock-transfected cells, consistent with effective inhibition of RNase L and preservation of host mRNA.

Together, these findings support a model in which RNase L activation by the E3 construct enhances transgene expression through selective depletion of host mRNA, thereby reducing competition for the ribosomal pool. Conversely, E3-NSs-L* suppresses RNase L activity, preserving host mRNA and limiting the availability of ribosomes for transgene expression.

### dsRNA-sensing pathway inhibition suppresses select antiviral cytokines while enhancing others

For therapeutic applications beyond immunotherapy, prolonged transgene expression is ideally achieved without triggering immune activation. While our data show that inhibition of dsRNA-sensing pathways can mitigate key saRNA-induced adverse effects, including cell death, translation shutdown, and host mRNA degradation, it remains unclear whether this strategy also attenuates cytokine secretion. To investigate this, we measured the concentrations of 13 antiviral cytokines in FLS culture supernatants using a bead-based multiplex immunoassay (Figure 4a and Supplemental Figure S4a-m). Transfection with native saRNA elicited a broad cytokine response. Inhibition of dsRNA-sensing pathways selectively suppressed production of certain cytokines: Both E3 and E3-NSs-L* significantly reduced interferon (IFN)-α and IFN-β, while E3-NSs-L* significantly reduced tumor necrosis factor (TNF). Notably, some production of cytokines was elevated: E3 transfection significantly increased macrophage chemoattractant protein (MCP)-1, and E3-NSs-L* significantly increased granulocyte-macrophage colony-stimulating factor (GM-CSF), relative to mock-transfected controls. These findings indicate that even strong inhibition of dsRNA-sensing pathways only partially attenuates the cytokine response to saRNA. Additional strategies may be needed to achieve fully non-immunostimulatory saRNA transfection suitable for non-immunotherapy applications.

**Figure 4.**
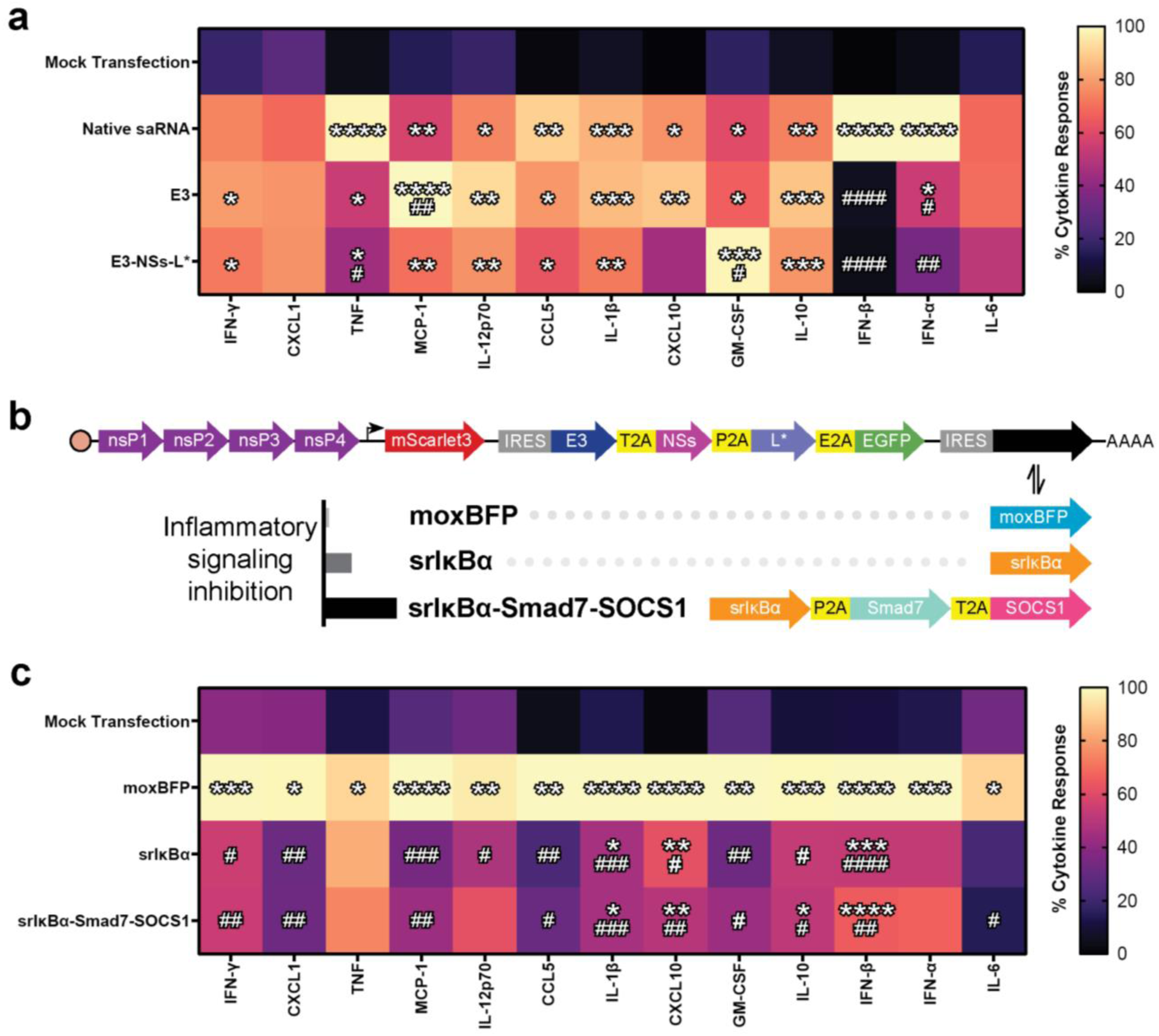
Inhibition of NF-κB signalling attenuates saRNA-induced secretion of antiviral cytokine production **a,** FLS were transfected with the saRNA constructs depicted in Fig. 1a, and antiviral cytokines in cell culture supernatant were quantified two days later using a bead-based immunoassay (n = 6 biological replicates). Inhibition of dsRNA-sensing pathways did not broadly suppress cytokine secretion, but significantly reduced saRNA-induced production of TNF, IFN-α and IFN-β. Cytokine levels were normalized to pre-transfection cell number (measured using BioTracker), scaled within each biological replicate (highest value set to 100%), and displayed as a heatmap of group means. Detailed plots for individual cytokines are shown in Supplemental Figure S4. Statistical significance was assessed using one-way repeated-measures ANOVA on unscaled data, with multiple comparisons controlled using the two-stage linear step-up procedure of Benjamini, Krieger, and Yekutieli (FDR = 5%). All cytokines passed significance thresholds for discovery. Treatment effects were analyzed using Tukey’s multiple comparisons test. *P < 0.05, **P < 0.01, ***P < 0.001, ****P < 0.0001 vs. mock transfection; ^#^P < 0.05, ^##^P < 0.01, ^###^P < 0.001 vs. native saRNA. The mock transfection control is shared with panel (c). **b,** Schematic of the moxBFP, srIκBα, and srIκBα-Smad7-SOCS1 saRNA constructs designed to inhibit inflammatory signalling. All constructs include dsRNA-sensing pathway inhibitors (vaccinia virus E3, Toscana virus NSs, and Theiler’s virus L*). The moxBFP construct, used as a control, lacks inflammatory signalling inhibitors. The srIκBα construct co-expresses srIκBα to block NF-κB signalling, representing moderate inflammatory signalling inhibition. The srIκBα-Smad7-SOCS1 construct co-expresses srIκBα, Smad7 and SOCS1 to additionally suppress TGF-β and IFN pathways, representing strong inhibition. The angled arrow denotes the subgenomic promotor. **c,** FLS were transfected with the saRNA constructs shown in (b), and antiviral cytokines were measured two days post-transfection (n = 6 biological replicates). Inhibition of NF-κB signalling with srIкBα broadly suppressed cytokine secretion. Data normalization and visualization were performed as described in (a), but scaling was applied independently to account for the different constructs tested. Detailed cytokine plots are shown in Supplemental Figure S5. Statistical significance was assessed using one-way repeated-measures ANOVA on unscaled data, with multiple comparisons controlled using the two-stage linear step-up procedure of Benjamini, Krieger, and Yekutieli (FDR = 5%). All cytokines met discovery thresholds. Treatment effects were assessed using Tukey’s test. *P < 0.05, **P < 0.01, ***P < 0.001, ****P < 0.0001 vs. mock transfection; ^#^P < 0.05, ^##^P < 0.01, ^###^P < 0.001, ^####^P < 0.0001 vs. moxBFP. The mock transfection control is shared panel (a). Acronyms: nsP, Venezuelan equine encephalitis virus non-structural protein; IRES, encephalomyocarditis virus internal ribosome entry site; moxBFP, monomeric oxidizing environment-optimized blue fluorescent protein; E3, Vaccinia virus E3 protein; NSs, Toscana virus non-structural NSs protein; L*, Theiler’s murine encephalomyelitis virus L* protein; T2A, Thosea asigna virus 2A peptide; P2A, porcine teschovirus-1 2A peptide, E2A, equine rhinitis A virus 2A peptide, srIкBα, super-repressor inhibitor of κBα; smad7, mothers against decapentaplegic homolog 7; SOCS1, suppressor of cytokine signalling 1; IFN-γ, interferon-γ; CXCL1, C-X-C motif chemokine ligand 1; TNF, tumor necrosis factor; MCP-1, monocyte chemoattractant protein-1; IL-12p70, interleukin-12; CCL5, chemokine ligand 5; IL-1β, interleukin-1β; CXCL10, C-X-C motif chemokine ligand 10; GM-CSF, granulocyte-macrophage colony-stimulating factor; IL-10, interleukin-10; IFN-β, interferon-β; IFN-α, interferon-α; IL-6, interleukin-6.

### Design of saRNA constructs for inhibiting multiple inflammatory signalling pathways

To further inhibit saRNA-induced cytokine release, we designed additional saRNA constructs that co-express cellular inhibitors of inflammatory signalling. In these constructs, dsRNA pathway inhibitors (vaccinia virus E3, Toscana virus NSs, Theiler’s virus L*) and EGFP were expressed from a single IRES (Figure 4b). A second IRES was then used to express varying combinations of inflammatory signalling inhibitors. The control construct, ‘moxBFP’, lacks inflammatory signalling inhibitors and instead expressed a blue fluorescent protein to confirm IRES functionality.

Given that activation of nuclear factor-кB (NF-κB) is a hallmark of viral infection and a master regulator of cytokine and chemokine induction,^63,64^ we prioritized its inhibition using super repressor inhibitor of κBα (srIκBα). This dominant active form of IκBα cannot be phosphorylated and thus resists ubiquitination and degradation, forming a stable cytoplasmic pool of IκBα that prevents NF-κB nuclear translocation and downstream signalling.^65,66^ We generated one construct expressing srIκBα alone (named ‘srIκBα’) and another expressing srIκBα in combination with Smad7 and suppressor of cytokine signalling 1 (SOCS1) using non-identical 2A peptides (named ‘srIκBα-Smad7-SOCS1’). Smad7 negatively regulates both transforming growth factor-β (TGF-β) and NF-κB signalling pathways,^67^ while SOCS1 inhibits type I, II, and III IFN pathways as well as NF-κB signalling.^68–70^

### Inhibiting NF-кB signalling broadly attenuates saRNA-induced antiviral cytokine secretion

Transfection with the moxBFP construct, which lacks inflammatory signalling inhibitors, led to a significant increase in the secretion of all measured cytokines (Figure 4c and Supplemental Figure S5a-m). Co-expression of srIκBα broadly reduced cytokine levels, significantly suppressing all cytokines except TNF, IFN-α and interleukin (IL)-6. The srIκBα-Smad7-SOCS1 construct did not further enhance suppression relative to srIκBα alone, with the exception of interleukin (IL)-6, which was significantly reduced only by srIκBα-Smad7-SOCS1. These results indicate that NF-κB is a major driver of saRNA-induced cytokine production, and that co-expression of srIκBα is an effective strategy to attenuate saRNA-induced inflammatory activation.

### Reduced antiviral gene expression and replicon activity observed with co-expression of inflammatory signalling inhibitors

To evaluate whether co-expression of anti-inflammatory proteins could attenuate saRNA-induced induction of antiviral transcripts, we performed a RT-qPCR array on mock-transfected cells and cells transfected with native saRNA, E3, or srIκBα-Smad7-SOCS1 (Supplemental Figure S6a). Transfection with native saRNA led to marked upregulation of several antiviral and proinflammatory mRNA transcripts, including *Adar*, *Isg20*, *Rigi*, *Ifih1*, *Tlr3*, *Eif2ak2*, *Zc3hav1* and *Il6*. In contrast, both the E3 and srIκBα-Smad7-SOCS1 constructs attenuated the induction of these transcripts, with srIκBα-Smad7-SOCS1 exhibiting broader suppression across the panel.

We also quantified *EGFP* transcript levels, present on all saRNA constructs, as a proxy for replicon amplification. FLS transfected with srIκBα-Smad7-SOCS1 exhibited significantly lower *EGFP* transcript levels than those transfected with native saRNA or E3 (Supplemental Figure S6b). Several factors may underlie this reduction in replicon activity, including the increased transcript length of srIκBα-Smad7-SOCS1 (∼40% longer than native saRNA or E3), reduced molar input at equal transfection mass, inhibition of RNase L (which may preserve host RNA and increase competition for translational and replication resources), and/or interference from a truncated transcript common to the moxBFP, srIκBα, and srIκBα-Smad7-SOCS1 constructs (Supplemental Figure S6c and d).

### srIκBα reduces cell number, whereas srIκBα-Smad7-SOCS1 preserves cell number and enhances transgene expression

We then examined the impact of inhibiting inflammatory signalling on cell number and transgene expression (Figure 5a and b). Interestingly, the srIκBα construct caused both immediate and sustained reductions in cell number, as quantified by BioTracker signal, whereas the srIκBα-Smad7-SOCS1 construct maintained cell number comparable to mock transfection throughout the experiment (Figure 5c). AUC analysis of the BioTracker data confirmed that srIκBα-Smad7-SOCS1 significantly mitigated the srIκBα-induced reduction in integrated BioTracker signal (Supplemental Figure S7a). Additional experiments using CellTag and calcein AM staining corroborated these findings, showing that srIκBα negatively impacted both cell number (Supplemental Figure S7b) and viability (Supplemental Figure S7c), effects that were prevented by co-expression of Smad7 and SOCS1.

**Fig. 5.**
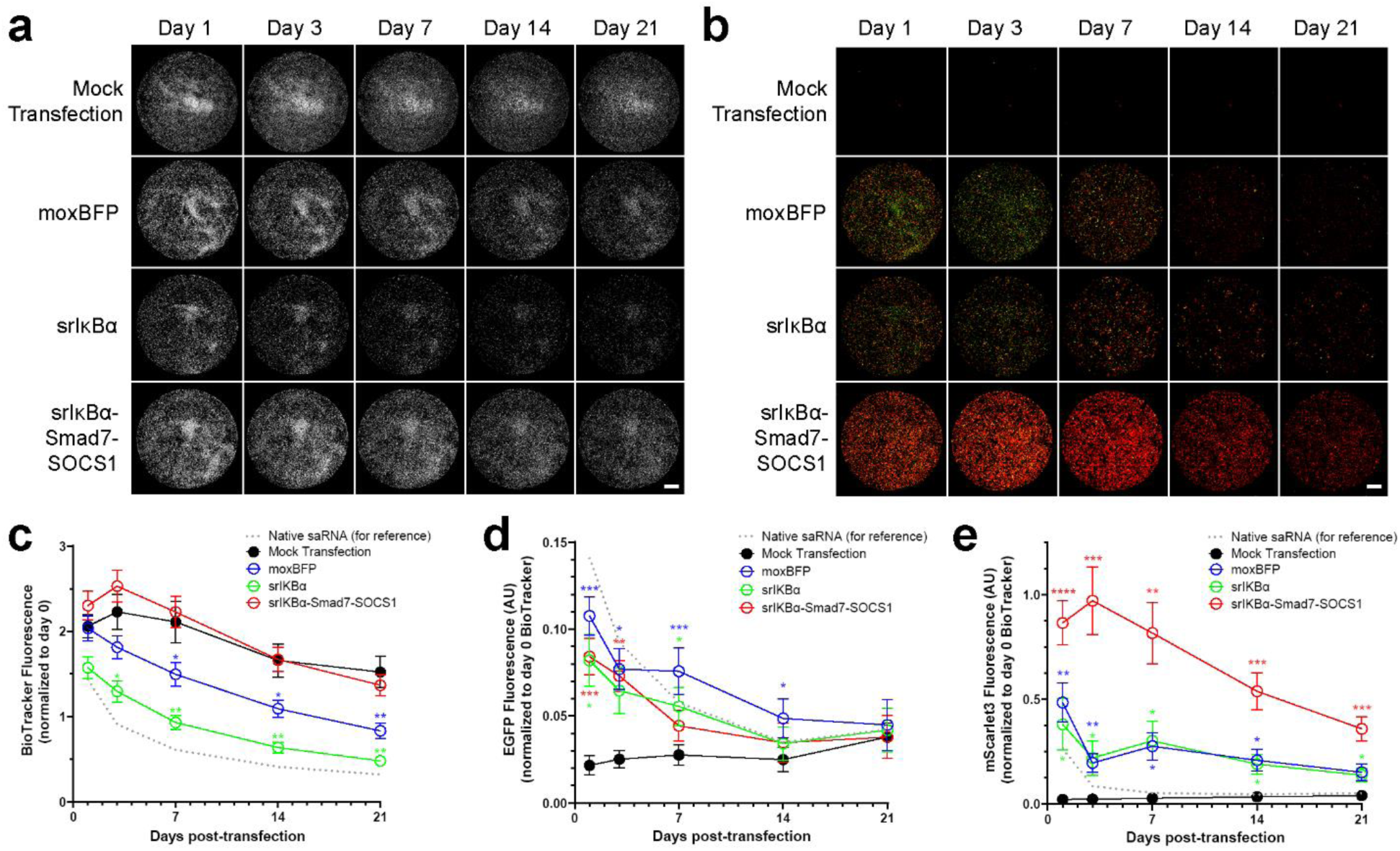
Differential effects of srIκBα and srIκBα-Smad7-SOCS1 on cell number and transgene expression. **a**, Representative images of BioTracker staining over 3 weeks in FLS transfected with different saRNA constructs. Scale bar = 5 mm. **b**, Representative images of EGFP (green) and mScarlet3 (red) expression over 3 weeks in FLS transfected with different saRNA constructs. Scale bar = 5 mm. **c**, Quantification of BioTracker fluorescence intensity over time (n = 11 biological replicates). srIκBα induces reduction in cell number, which is prevented by srIκBα-Smad7-SOCS1. Statistical significance of treatment effects at each time point compared to mock transfection was determined by two-way RM ANOVA with Greenhouse–Geisser correction and Dunnett’s multiple comparisons test. **d**, Quantification of EGFP fluorescence intensity over time (n = 11 biological replicates). All constructs showed low levels of EGFP expression. Statistical significance of treatment effects at each time point compared to mock transfection was determined by two-way RM ANOVA with Greenhouse–Geisser correction and Dunnett’s multiple comparisons test. **e**, Quantification of mScarlet3 fluorescence intensity over time (n = 11 biological replicates). The srIκBα-Smad7-SOCS1 produced 2-5 times more mScarlet3 fluorescence than either moxBFP or srIκBα constructs. Statistical significance of treatment effects at each time point compared to mock transfection was determined by two-way RM ANOVA with Greenhouse–Geisser correction and Dunnett’s multiple comparisons test. saRNA constructs used are shown in Figure 4b. For reference, a dotted line shows the responses to native saRNA in panels c–f. For all statistical reporting, *P < 0.05, **P < 0.01, ***P < 0.001 and ****P < 0.0001. Data were normalized to cell number (BioTracker intensity) on day 0, prior to transfection. The mock transfection control data used in this figure is also presented in Figure 1d–f. Data are presented as mean ± SEM.

All three constructs produced low levels of EGFP expression (Figure 5d), which was expected given EGFP’s position as the fourth protein in a quad-cistronic 2A element (Figure 4b). Previous studies have demonstrated that protein expression decreases significantly for downstream genes in such configurations.^41^ Notably, the srIκBα-Smad7-SOCS1 construct produced 2-to 5-fold higher mScarlet3 fluorescence than either the moxBFP or srIκBα constructs (Figure 5e). This indicates that the srIκBα-Smad7-SOCS1 construct supports robust cap-dependent transgene expression despite its multi-cistronic configuration.

### Smad7 and SOCS1 co-expression prevents srIκBα-induced alterations in translational control

We next explored how srIκBα and srIκBα-Smad7-SOCS1 modulate translational regulation. Interestingly, srIκBα expression led to reductions in eIF2α phosphorylation, a change that was prevented by co-expression of Smad7 and SOCS1 (Figure 6a). While neither construct affected total eIF2α levels or eIF4E phosphorylation compared to the moxBFP construct (Figure 6b and c), srIκBα caused a significant reduction in total eIF4E levels (Figure 6d). Given the critical role of eIF4E in cap-dependent translation, its depletion likely contributes to the poor transgene expression observed with srIκBα (Figure 5e). These effects, combined with srIκBα-induced declines in cell number and viability (Figure 5c and Supplemental Figure S7a–c), highlight the dual impact of srIκBα on translational machinery and cellular health.

**Fig. 6.**
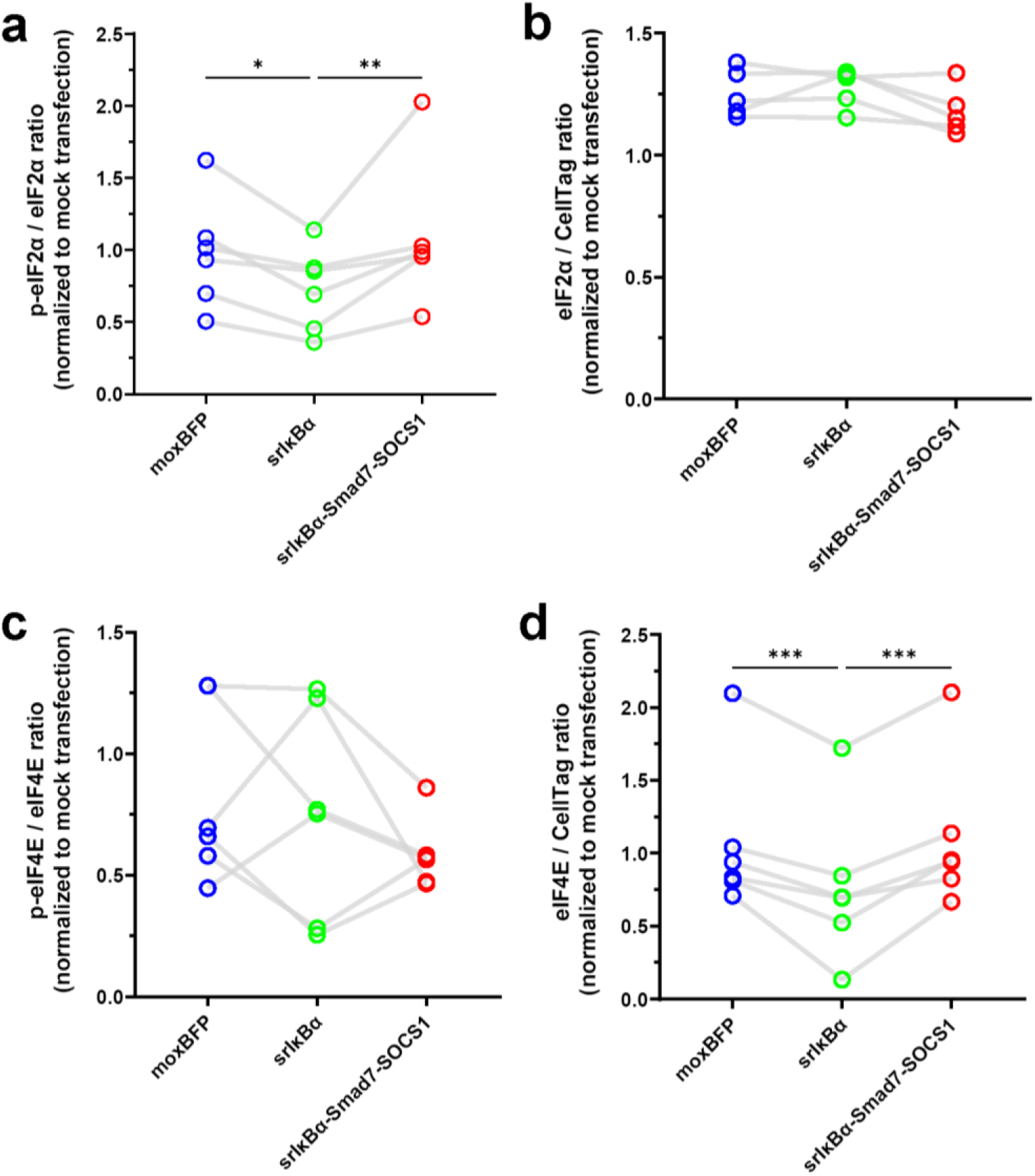
srIκBα reduces eIF2α phosphorylation and total eIF4E levels, effects reversed by co-expression of Smad7 and SOCS1. In-cell western assays were performed 2 days post-transfection. Data are presented as fold change relative to mock-transfected cells. **a,** Phosphorylation of eIF2α is significantly reduced by srIκBα, and this reduction is reversed by co-expression of Smad7 and SOCS1 (n = 6 biological replicates). Statistical significance was determined by one-way RM ANOVA and Holm-Šídák’s multiple comparisons test to compare all groups. **b,** Total eIF2α levels are not significantly affected by srIκBα or srIκBα-Smad7-SOCS1 (n = 6 biological replicates). One-way RM ANOVA revealed no significant differences among groups. F(2,8)=3.683, P=0.0735. **c,** Phosphorylation of eIF4E is not significantly affected by srIκBα or srIκBα-Smad7-SOCS1 (n = 6 biological replicates). One way RM ANOVA revealed no significant difference among groups. F(2,10)=1.336, P=0.3059. **d,** Total eIF4E levels are significantly reduced by srIκBα, an effect reversed by co-expression of Smad7 and SOCS1 (n = 6 biological replicates). Statistical significance was determined by one-way RM ANOVA and Holm-Šídák’s multiple comparisons test to compare all groups. saRNA constructs are described in Figure 4b. Connecting lines indicate responses from the same biological replicate. For all statistical reporting, *P < 0.05, **P < 0.01 and ***P < 0.001. Mock transfection data used for normalization are the same as in Fig. 3.

Importantly, Smad7 and SOCS1 co-expression preserved eIF4E levels, counteracting the disruption of cap-dependent translation caused by srIκBα.

### Prolonged moxBFP transfection does not activate FLS, while srIκBα reduces basal activation

Fibroblast activation protein-α (FAP-α) is a serine protease expressed on the surface of FLS that contributes to extracellular matrix degradation and tissue remodeling.^71,72^ Its expression is minimal in normal adult FLS but increases significantly following inflammatory activation, such as in rheumatoid and osteoarthritis.^73,74^ Since viral infections can induce fibroblast activation,^75,76^ we hypothesized that prolonged saRNA transfection might similarly trigger FLS activation. To test this, we measured FAP-α levels using an in-cell western assay after 11 days of transfection. FLS transfected with moxBFP showed FAP-α levels comparable to those of mock-transfected cells, whereas srIκBα and srIκBα-Smad7-SOCS1 constructs resulted in significantly lower FAP-α levels compared to mock transfection (Figure 7a and b). These findings indicate that long-term saRNA transfection does not induce FLS activation, as measured by FAP-α levels, potentially alleviating concerns that saRNA might drive pathological fibroblast responses.

**Figure 7.**
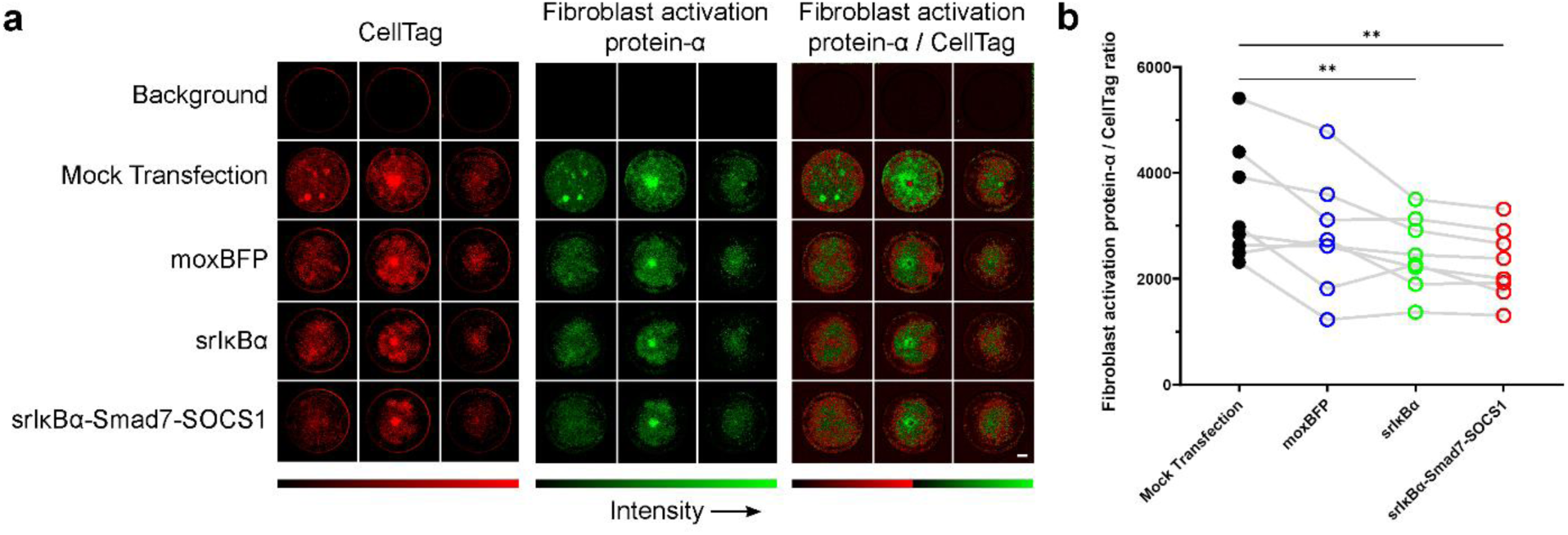
Prolonged transfection with *srIκBα or srIκBα-Smad7-SOCS1 significantly reduces basal fibroblast activation factor-α (FAP-α) levels* a, Representative in-cell western images showing FAP-α expression. Columns show different biological replicates, and rows show different treatments. The montage on the right shows FAP-α signal normalized to CellTag signal (FAP-α/CellTag). **b,** In-cell western assay of FAP-α expression (n = 8 biological replicates). Both srIκBα and srIκBα-Smad7-SOCS1 significantly reduce FAP-α levels compared to mock transfection, while moxBFP does not differ significantly from mock. Statistical significance was determined by one-way RM ANOVA with Greenhouse–Geisser correction and Dunnett’s multiple comparisons test to compare groups to mock transfection. Connecting lines indicate responses from the same biological replicate. **P<0.01.

### External control of transgene expression using a small-molecule antiviral

The srIκBα-Smad7-SOCS1 construct represents a promising immune-evasive saRNA platform for transient gene therapy applications. It includes the E3-NSs-L* polyprotein, which protects against saRNA-induced cytotoxicity (Figure 1f; Figure 3—figure supplement 1a–b; Figure 2b and d), translation shutdown (Figure 1e; Figure 3a), and host mRNA degradation (Figure 3—figure supplement 1b–c). In parallel, co-expression of srIκBα attenuates saRNA-induced cytokine responses (Figure 4c), while Smad7 and SOCS1 mitigate srIκBα-associated reductions in cell viability (Figure 5c; Supplemental Figure 7a–c) and eIF4E levels (Figure 6d). These combined features enable sustained cap-dependent transgene expression (Figure 5e) without triggering fibroblast activation (Figure 7b).

To improve safety and flexibility, we next investigated whether transgene expression could be externally regulated using a small-molecule antiviral. ML336 is a potent inhibitor of the VEEV RdRp used by the saRNA replicon.^45^ To evaluate its efficacy in modulating transgene output, we performed a concentration-response assay by adding ML336 to the culture medium during transfection. ML336 suppressed mScarlet3 expression in a concentration-dependent manner (Figure 8a), with an IC_50_ of 8.5 nM, demonstrating effective and tunable external control of saRNA-driven expression.

**Fig. 8.**
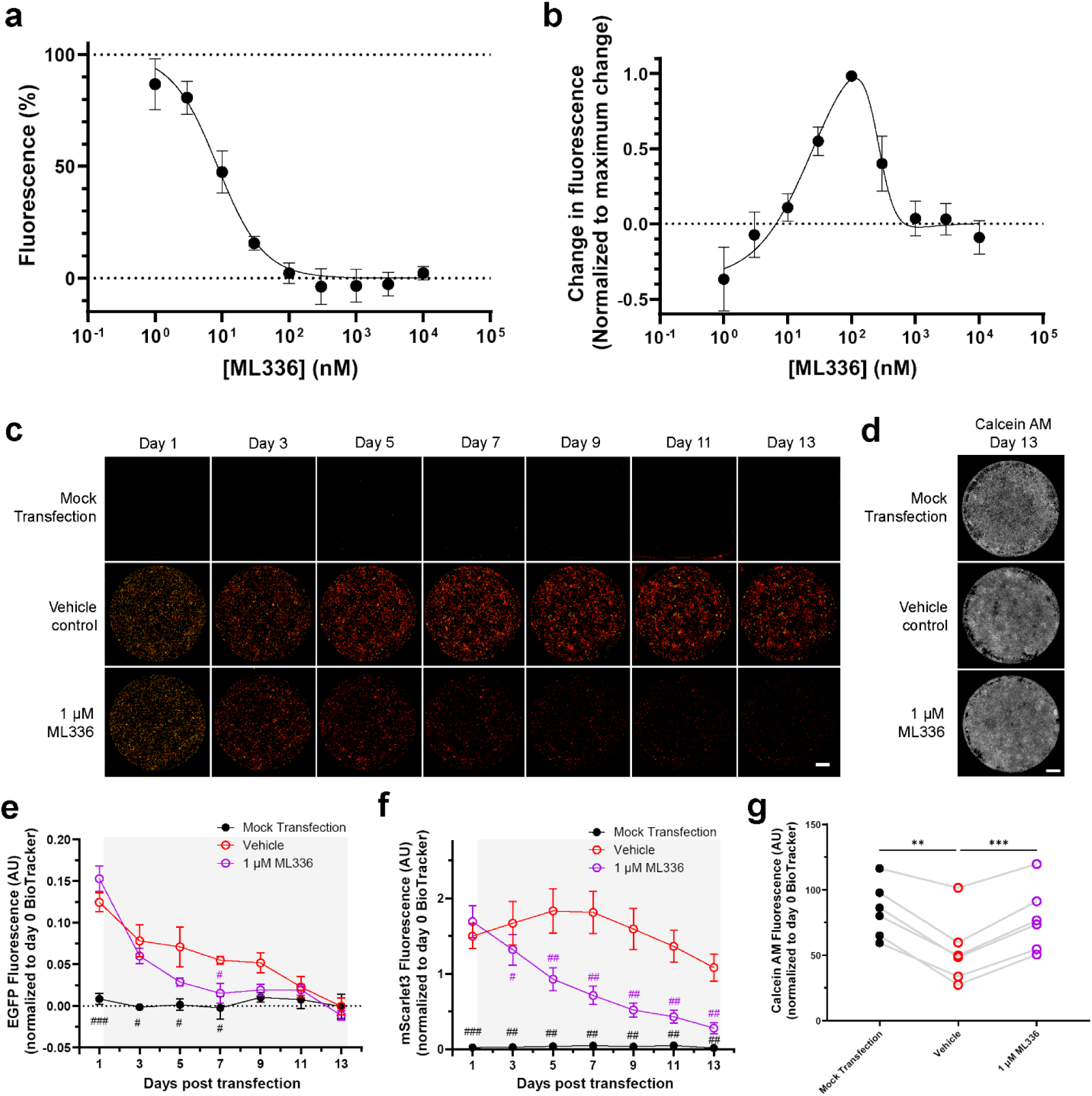
ML336 enables reversible external control of transgene expression from the srIκBα-Smad7-**SOCS1 construct a,** FLS were transfected with the srIκBα-Smad7-SOCS1 construct and treated with increasing concentrations of ML336. mScarlet3 fluorescence was measured two days post-transfection and normalized to vehicle-treated controls (n = 5 biological replicates). The concentration-response curve was fit using a variable-slope sigmoidal model, yielding an IC_50_ of 8.5 nM (95% CI: 6.1–11.9 nM). **b,** ML336 was removed from the media on day 2 post-transfection, and mScarlet3 fluorescence was measured on day 3 (n = 5 biological replicates). Fluorescence changes were normalized to the maximum response within each biological replicate and fit to a bell-shaped concentration-response curve. ML336 removal at concentrations between 10 nM and 1 μM led to disinhibition of the RdRp and increased mScarlet3 expression. **c,** Representative images of EGFP (green) and mScarlet3 (red) expression over 13 days in FLS transfected with srIкBα-smad7-SOCS1. Cultures were treated with vehicle or 1 μM ML336, starting 1 day post-transfection. Scale bar = 5 mm. **d,** Representative images of calcein AM staining on day 13 post-transfection. Scale bar = 5 mm. **e,** Quantification of EGFP fluorescence intensity in srIκBα-Smad7-SOCS1-transfected FLS (n = 6 biological replicates). Cultures were treated with vehicle or 1 μM ML336 (treatment period indicated by shading). EGFP fluorescence was significantly lower in ML336-treated cultures compared to vehicle-treated cultures on day 7. Statistical significance relative to vehicle-treated cells was determined by two-way RM ANOVA with Greenhouse–Geisser correction and Dunnett’s multiple comparisons test. ^#^P < 0.05, ^###^P < 0.01. **f,** Quantification of mScarlet3 fluorescence intensity in srIκBα-Smad7-SOCS1-transfected FLS (n = 6 biological replicates). Cultures were treated with vehicle or 1 μM ML336 (treatment period indicated by shading). mScarlet3 fluorescence was significantly lower in ML336-treated cultures compared to vehicle-treated cultures beginning on day 3. Statistical significance relative to vehicle-treated cells was determined by two-way RM ANOVA with Greenhouse–Geisser correction and Dunnett’s multiple comparisons test. ^#^P < 0.05, ^##^P < 0.01, ^###^P < 0.001. **g,** Quantification of calcein AM staining on day 13 post-transfection, following 12 days of vehicle or ML336 treatment (n = 6 biological replicates). ML336-treated cultures showed no significant difference in calcein AM signal compared to mock transfection, while vehicle-treated cultures exhibited significantly lower calcein AM signals compared to both mock-transfected and ML336-treated cultures. Connecting lines indicate responses from the same biological replicate. Statistical significance was assessed by one-way RM ANOVA with Greenhouse–Geisser correction and Tukey’s multiple comparisons test. **P<0.01, ***P<0.001. All fluorescence data were normalized to BioTracker intensity on day 0 and are presented as mean ± SEM.

We then assessed whether ML336 could be used to eliminate the replicon, providing a mechanism to terminate transgene expression after therapeutic goals have been met. ML336 was removed from the cell culture medium two-days post transfection, and mScarlet3 fluorescence was measured on the following day. At intermediate concentrations (10 nM–1 μM), where ML336 inhibits subgenomic RNA synthesis without completely blocking replication, mScarlet3 expression partially recovered, with maximal recovery at 100 nM (Figure 8b). In contrast, no recovery was observed at concentrations above 1 μM, suggesting that high ML336 levels inhibit genomic saRNA replication and result in replicon clearance.

To assess long-term control of transgene expression, FLS were transfected with the srIκBα-Smad7-SOCS1 construct and treated with either 1 μM ML336 or vehicle one day post-transfection. EGFP and mScarlet3 fluorescence were monitored over 13 days (Figure 8c), followed by calcein AM staining to assess cell viability (Figure 8d). ML336 treatment significantly reduced EGFP and mScarlet3 expression over time (Figure 8e and f), demonstrating its utility for sustained external control of transgene expression. Importantly, prolonged ML336 treatment did not impair cell viability relative to mock-transfected controls (Figure 8g). In contrast, vehicle-treated cultures exhibited reduced calcein AM staining. This reduction may be due to decreased cell proliferation over the course of the experiment, which is consistent with our observations that saRNA reduces eIF4E phosphorylation (Figure 3c), a change associated with impaired cell growth and proliferation.^77^

## DISCUSSION

Although saRNA is known to disrupt translational control, the precise mechanisms and downstream effects remain incompletely characterized. saRNA-induced translation shutdown is thought to be mediated by phosphorylation of eIF2α,^21,78,79^ mediated by activation of PKR in response to dsRNA replicative intermediates.^7,21^ Our findings support this model and further show that saRNA reduces eIF4E phosphorylation—a phenomenon previously observed with alphavirus replicons, though its mechanism is not yet fully understood.^80,81^ Reduced eIF4E phosphorylation suppresses cap-dependent translation initiation,^82^ thereby favouring cap-independent translation.^83^ Together, eIF2α phosphorylation and reduced eIF4E activity likely explain why IRES-driven transgenes achieve higher expression than their cap-dependent counterparts when using native saRNA.^26,84^

In addition to disrupting translation initiation, saRNA replication induces host translation shutoff through activation of RNase L. Many viruses exploit host shutoff—the global suppression of host gene expression, as a strategy to redirect cellular resources toward their replication.^85^ saRNA appears to co-opt this mechanism: RNase L activation depletes host mRNAs without impairing the translation machinery.^55^ However, saRNA transcripts, are largely protected from degradation due to their localization within viral replication organelles known as spherules.^3,78^

RNase L activity extends beyond transcript degradation. It also cleaves actively translated mRNAs, leading to ribosome stalling at 3’ ends, ribosome collisions,^86,87^ activation of the ribotoxic stress response,^88^ and ultimately, apoptosis.^86,89^ PKR likewise contributes to saRNA-induced cytotoxicity and can induce apoptosis independently of eIF2α phosphorylation.^90^ For example, microinjected dsRNA induces apoptosis in a PKR-dependent manner in HeLa cells, with RIG-I and MDA-5 being dispensable.^91^ Additional studies further support PKR as a central mediator of apoptosis in response to cytosolic dsRNA.^92–94^

These cytotoxic effects are exemplified by the E3 construct, which alleviates translation shutdown but does not inhibit RNase L activation. The resulting depletion in host transcripts reduces ribosomal competition, enhancing saRNA transgene translation—but at the cost of long-term cytotoxicity. This dual-edged effect may make the E3 construct well suited for immunotherapy applications, such as vaccines or cancer therapy, where robust transgene expression and immunogenic cell death are advantageous.^95,96^ In contrast, applications requiring sustained cell viability demand strategies that mitigate both saRNA-induced translation shutdown and RNase L–mediated host shutoff. The E3-NSs-L* construct, which combines broad inhibition of dsRNA-sensing pathways with targeted suppression of PKR and RNase L, effectively protects against the intrinsic cytotoxicity associated with saRNA replication.

While we found that inhibiting dsRNA-sensing pathways was an effective means to reduce saRNA-induced cytotoxicity, it was not effective at preventing saRNA-induced cytokine responses. This agrees with prior reports showing that dsRNA activates NF-κB even in the absence of PKR and RNase L,^97^ suggesting that alternative pathways contribute to NF-κB activation. One such mechanism may involve eIF4E, whose activity is governed by its phosphorylation status and serves as a rate-limiting step in cap-dependent protein synthesis.^52^ In this study, we found that saRNA transfection induces a reduction in eIF4E phosphorylation. Reduced eIF4E phosphorylation has been shown to decrease the levels of the short-lived IκBα protein,^98^ an inhibitor of NF-κB that sequesters it in the cytoplasm and prevents nuclear translocation. Decreased IκBα levels thus allow NF-κB to translocate to the nucleus, initiating cytokine expression and inflammatory responses.^99^

Importantly, while NF-κB is best known for promoting inflammatory cytokine production, it also supports cell survival by inducing anti-apoptotic genes, though its effects vary by tissue type and biological context.^100^ In FLS, overexpression of srIκBα reduces cytokine production but increases sensitivity to TNF-induced apoptosis,^101,102^ consistent with our findings that saRNA encoding srIκBα lowered both cytokine responses and cell viability. This phenomenon is not unique to FLS; NF-κB inhibition with srIκBα also renders liver hepatocytes more susceptible to pro-apoptotic stimuli,^103^ raising safety concerns given that RNA delivery systems such as lipid nanoparticles (LNPs) tend to accumulate in the liver.^104^ Therefore, when designing saRNA constructs for sustained, non-immunostimulatory transgene expression in contexts where apoptosis is undesirable, the use of srIκBα to suppress cytokine responses may require co-expression of anti-apoptotic proteins to offset the loss of NF-κB-mediated survival signalling. In our case, co-expressing Smad7 and SOCS1 restored FLS viability in the presence of srIκBα. While we did not assess the individual contributions of each protein, both are known to have anti-apoptotic properties in certain contexts. Smad7 can inhibit TGF-β-induced apoptosis,^105,106^ while SOCS1 can suppress apoptosis triggered by IFN-α, IFN-β,^107^ IFN-γ,^108^ and TNF-α.^109–111^ These anti-apoptotic effects likely enable Smad7 and SOCS1 to mitigate srIκBα-induced reductions in FLS cell viability.

We also found that co-expression of Smad7 and SOCS1 significantly enhanced cap-dependent transgene expression. Although we did not distinguish the contributions of each protein to this effect, it is likely mediated by SOCS1. Previous studies have shown that ruxolitinib, a small-molecule inhibitor of Janus kinase (JAK) 1 and JAK2,^112^ increases saRNA transgene expression when added to saRNA polyplexes.^18^ SOCS1 is an endogenous inhibitor of JAK1 and JAK2;^113^ therefore, SOCS1 offers a genetically encoded alternative to the small-molecule approach of ruxolitinib for enhancing saRNA expression.

Furthermore, this study presents a straightforward method for controlling saRNA activity. ML336, a VEEV RdRp inhibitor, blocks the synthesis of positive-sense genomic, negative-sense template, and subgenomic RNAs of VEEV without affecting host cellular RNA transcription.^114^ It has an *in vitro* effective concentration 50 (EC_50_) of 0.02-0.04 μM, and a 50 mg/kg dose offers 100% protection in a lethal VEEV mouse infection model, with no observed toxicity.^115^ Thus, ML336 and other RdRp inhibitors could provide a practical means to deactivate saRNA—either to terminate transgene expression once therapeutic objectives are met or as a safety measure to reduce the risks associated with prolonged innate immune suppression and other potential side effects, including adaptive immune responses to the VEEV replicon or viral innate immune inhibitory proteins.

Preclinical and clinical data show that saRNA vaccines often trigger an intense innate immune response that can hinder antigen expression.^8,18,21^ Additionally, saRNA vaccines can induce significant inflammation,^116,117^ which is increasingly recognized as problematic, as common adverse events include pain, headache, tenderness, arthralgia, fever, and chills.^118,119^ The approaches described in this study, which reduces the inherent immunostimulatory effects of saRNA replication, could potentially be applied to future saRNA vaccine designs to improve antigen expression and reduce reactogenicity.

The approach in this study also underscores the versatility of the saRNA platform, which can efficiently integrate multiple transgenes and genetic elements. Our largest construct included the VEEV RdRP, two IRES sequences, five 2A peptides, and eight transgenes within a 16.5-kilobase saRNA. Leveraging this capacity for multiple transgenes, we encoded diverse inhibitors of dsRNA-sensing and inflammatory signalling pathways directly within saRNA achieving durable and controllable transgene expression with minimal cytotoxicity and immunostimulation. This approach eliminates the need for exogenous immune suppressants—a common requirement in other studies utilizing saRNA for non-immunotherapy applications.^9,20,22,26,27^ By localizing immune suppression to transfected cells, this approach reduces treatment complexity and minimizes the risk of systemic immunosuppression.

Expressing multiple transgenes may also be a useful strategy for developing novel therapeutics. The NF-κB, TGF-β, and JAK/signal transducer and activator of transcription (STAT) pathways are central to the progression of osteoarthritis.^120–122^ The srIκBα-Smad7-SOCS1 saRNA construct provides a novel strategy to simultaneously target these pathways: srIκBα inhibits NF-κB, Smad7 blocks TGF-β, and SOCS1 suppresses the JAK/STAT pathway. Each of these proteins has been independently studied as a treatment for osteoarthritis,^123–125^ and their combination within a single construct could represent a synergistic strategy to enhance therapeutic efficacy. While arthritogenic alphaviruses such as chikungunya virus and Ross river virus can drive inflammatory responses in FLS, including upregulation of secreted proteolytic molecules such as matrix metalloproteases,^126,127^ the lack of FAP-α upregulation in transfected FLS provides some evidence that saRNA transfection does not trigger comparable responses. Moreover, the reduction in basal FAP-α levels observed with the srIκBα and srIκBα-Smad7-SOCS1 constructs raises the possibility that saRNA could serve as a tool for suppressing fibroblast activation in inflammatory settings. This approach could hold therapeutic potential in arthritis treatment,^128^ warranting further investigation in disease models.

Our findings present a fully saRNA-based strategy that addresses a central limitation of the platform: innate immune activation driven by replicon activity. By encoding proteins that inhibit both dsRNA-sensing pathways and NF-κB signalling directly within the saRNA—expressed via cap-independent translation— we achieved broad suppression of innate immune responses. This approach enabled sustained transgene expression with minimal cytotoxicity and antiviral cytokine release, eliminating the need for exogenous immunosuppressants. Furthermore, the use of ML336 enables on-demand regulation or reversal of transgene expression, enhancing both the safety and versatility of this approach. Given the established success of saRNA in preclinical models^6,18,21,26^ and approved human vaccines^119,129^ we anticipate that this immune-evasive saRNA strategy will be readily translatable to future *in vivo* studies.

However, the inclusion of multiple viral proteins introduces additional challenges. *In vivo*, saRNA constructs must contend not only with intrinsic cytotoxic mechanisms but also with adaptive immune responses that drive extrinsic cytotoxicity. Co-expression of several viral proteins, along with the VEEV RdRp, increases the likelihood that viral peptides will be presented on major histocompatibility complex class I molecules. This, in turn, is expected to promote recognition and elimination of saRNA-transfected cells by cytotoxic T lymphocytes, thereby limiting therapeutic durability. Overcoming both intrinsic and immune cell-mediated cytotoxicity will be essential for achieving long-term persistence of saRNA-transfected cells and realizing the platform’s full therapeutic potential.

In addition to characterizing the fate of saRNA-transfected cells *in vivo*, future studies should evaluate the biodistribution of immune-evasive saRNA. While mRNA-LNPs typically target the liver and induce strong hepatic expression, saRNA-LNPs, despite similar biodistribution, show little to no expression in the liver and instead exhibit significant expression in the spleen, lungs, and muscle^117,130^. This is likely due to differences in innate immune responses to mRNA versus the cytosolic dsRNA intermediates generated by saRNA. Accordingly, the immune-evasive strategy described here, which disrupts innate immune sensing of dsRNA, is likely to influence the biodistribution and expression profile of saRNA constructs, potentially resulting in unanticipated secondary effects. Further investigation into the *in vivo* fate and biodistribution of these constructs will be essential for optimizing their safety and efficacy across diverse clinical applications. Ultimately, continued refinement of immune-evasive saRNA constructs and delivery strategies will be crucial for achieving safe, sustained, reversible, and clinically viable saRNA-based gene therapies.

## MATERIALS & METHODS

### Experimental Design

This study was designed to develop and evaluate saRNA constructs capable of suppressing multiple innate immune pathways—specifically dsRNA-sensing and inflammatory signalling—to mitigate saRNA-induced immunotoxicities such as translation shutdown, cytotoxicity, mRNA degradation, and cytokine release. The overarching goal was to enable sustained transgene expression suitable for non-immunotherapeutic applications of saRNA, without the need for exogenous immunosuppressants.

To this end, we engineered saRNA constructs that co-express multiple immune-modulatory proteins, including viral inhibitors of dsRNA-sensing pathways and cellular inhibitors of inflammatory signalling. In addition, constructs were designed to enable simultaneous monitoring of cap-dependent and cap-independent translation via distinct fluorescent protein reporters.

The experimental workflow involved molecular cloning of the saRNA constructs, *in vitro* transcription, and transfection into primary mouse FLS. Transfected cells were assessed using fluorescence-based microplate assays to evaluate cell number, viability, phosphatidylserine exposure, and reporter protein expression. Further analyses comprised in-cell western assays for translational regulators and FLS activation markers, SUnSET assay for protein synthesis rates, capillary electrophoresis for rRNA integrity, and bead-based immunoassays for cytokine secretion. Finally, the potential for small-molecule regulation of transgene expression was tested using an antiviral compound that inhibits the VEEV RdRp.

### Plasmid cloning and construct design

The commercially available TagGFP2 E3L Simplicon vector (SCR725, Merck) served as the saRNA backbone in this study. After removal of the TagGFP2-IRES-E3 sequence, various constructs were generated by combining different elements using restriction digests (FastDigest, Thermo Scientific), overlap extension PCR (Phusion, Thermo Scientific), and HiFi assembly (NEBuilder, New England Biolabs).

Genetic elements were sourced as follows: mScarlet3 from pDx_mScarlet3 (a gift from Dorus Gadella [Addgene plasmid #189754]),^131^ IRES-EGFP from pIRES2-EGFP (Clontech), and IRES-E3 and IRES-PuroR (puromycin resistance) from the original TagGFP2 E3L Simplicon vector.

Custom gene synthesis (GeneArt, Thermo Scientific) was used to produce IRES-moxBFP, IRES-E3-T2A-NSs-P2A-L*-E2A-mEGFP, and IRES-srIкBα-P2A-Smad7-T2A-SOCS1 sequences. The IRES sequence was derived from pIRES2-EGFP (Clontech). Other sequences were obtained from public databases: moxBFP (FPbase ID: SSTDU),^132^ mEGFP (FPbase ID: QKFJN), E3 (UniProt: P21605-1), NSs (UniProt: P21699), L* (UniProt: P0DJX4), Smad7 (UniProt: O35253-1), and SOCS1 (UniProt: O35716). srIкBα was engineered from IкBα (UniProt: Q9Z1E3) by substituting serines 32 and 36 with alanines. All synthesized sequences were codon-optimized for mouse expression (GenSmart, GenScript).

Plasmids were cloned in DH5α competent *Escherichia coli* (High Efficiency, New England Biolabs) and purified by maxiprep (PureLink HiPure, Invitrogen). All plasmid sequences were verified using nanopore whole plasmid sequencing (Plasmidsaurus).

### RNA synthesis

Plasmids were linearized with XbaI (FastDigest, Thermo Scientific) at 37 °C for 3 hours. The linear plasmids were purified by phenol-chloroform extraction followed by sodium acetate-ethanol precipitation. Uncapped RNA was synthesized *in vitro* using the T7 RiboMAX Large Scale RNA Production System (Promega) at 37 °C for 2 hours. After purification by ammonium acetate precipitation, the RNA was denatured by heating to 65 °C for 5 minutes before rapidly cooling on ice. Cap-1 structures were then generated using the Vaccinia Capping System in conjunction with mRNA cap 2’-O-methyltransferase (both from New England Biolabs) for 45 minutes at 37 °C. Following another round of ammonium acetate precipitation, the RNA was treated with Antarctic phosphatase (New England Biolabs) for 30 minutes at 37 °C. After a final ammonium acetate precipitation, RNA was resuspended in THE RNA Storage Solution (Invitrogen). RNA concentration was quantified using an N60 NanoPhotometer (Implen) and adjusted to a final concentration of 0.5 μg/μL. RNA was aliquoted and stored at –80 °C until further use.

The integrity of the *in vitro* transcribed RNA was assessed by denaturing agarose gel electrophoresis (Supplemental Figure S6c). Constructs designed to inhibit dsRNA-sensing pathways (native saRNA, E3, and E3-NSs-L*) produced a single band at the expected size. In contrast, constructs targeting inflammatory signalling (moxBFP, srIκBα, and srIκBα-Smad7-SOCS1) displayed two bands: one corresponding to the full-length transcript, and a smaller, low-intensity band of consistent size across all three constructs (Supplemental Figure S6d). The presence of this truncated species likely reflects premature termination at a cryptic terminator introduced during construct assembly.

### Denaturing RNA gel electrophoresis

*In vitro* transcribed RNA (1 μg) was mixed with 5 μl of NorthernMax-Gly Sample Loading Dye (Invitrogen) and 5 μl of nuclease-free water for glyoxal-based denaturation. The RNA ladder (RiboRuler High Range RNA Ladder, Thermo Scientific) was prepared by combining 2 μl of ladder, 3 μl of water, and 5 μl of loading dye. A loading dye only control was also prepared using 5 μl of water and 5 μl of loading dye. All samples were incubated at 50 °C for 30 minutes, then cooled on ice.^133,134^

Samples were then loaded onto a 0.8% agarose gel prepared with diethyl pyrocarbonate (DEPC)-treated electrophoresis buffer consisting of 100 mM PIPES, 300 mM Bis-Tris, and 10 mM EDTA. Electrophoresis was performed at 5 V/cm, and gels were imaged using an Odyssey Fc imager (LI-COR) with acquisition in the 600 nm channel over 10 minutes.

### Mouse primary FLS culture

All mice used in this study were 5-to 12-week-old wildtype C57BL/6J mice (Envigo, Bicester, UK). Both male and female mice were included, though sex differences in response to saRNA transfection were not analyzed. Mice were housed in groups of up to five in a temperature-controlled room (21 °C), with bedding, a red shelter, and enrichment materials provided. They were maintained on a 12-hour light/dark cycle, with food and water available ad libitum. Mice were euthanized by exposure to CO_2_ gas in a rising concentration, and death confirmed by dislocation of the atlanto-occipital joint by applying pressure to the side of the neck using thumb and forefinger, procedures in accordance with the Animals (Scientific Procedures) Act 1986 Amendment Regulations 2012.

We cultured FLS from mouse knee joints following established protocols,^135^ with slight modifications. In brief, knee joints were exposed by cutting through the overlying skin with dissecting scissors. The patellar tendon was then grasped near the patella with Dumont tweezers, and spring scissors were used to sever the patellar tendon along with surrounding tissues. Finally, the patella was excised by cutting the quadriceps tendon, and excess muscle and tendons were carefully trimmed away.

Both patellae from each animal were collected into a microcentrifuge tube containing ice-cold sterile phosphate buffered saline (PBS), then transferred to a 24-well tissue culture plate (Costar) with FLS culture media. The media consisted of DMEM/F-12 (Invitrogen) supplemented with 25% fetal bovine serum (Sigma), 1X GlutaMAX (Gibco), and 100 μg/ml Primocin (InvivoGen).

The patellae were maintained in a humidified incubator at 37 °C with 5% CO2, with media changes every 1-2 days. FLS outgrowth was observed, and after approximately 10 days, when cells reached 70% confluency, the patellae were transferred to new wells for further FLS collection. Cells attached to the original wells were passaged using 0.1% trypsin-EDTA (Sigma) into a single well of a 6-well plate (Costar). Cells were later expanded into T25 flasks (BioLite, Thermo Scientific or CELLSTAR, Greiner Bio-One) and passaged upon reaching confluency. After the second passage, Versene solution (Gibco) was used for cell dissociation.

FLS were used between passages 3 and 8. In most experiments, cells from the same animal were divided and plated onto 6-well or 24-well plates, allowing for parallel manipulations with matched conditions. Occasionally, when cell number was low, cells from different animals were pooled before plating. Once plated for experiments, media was changed every 2–3 days.

### Immunocytochemistry and confocal microscopy

Cells were plated on poly-D-lysine–coated glass-bottom 35 mm dishes (P35GC-1.5-14-C, MatTek) and fixed with 4% paraformaldehyde for 10 minutes at room temperature without permeabilization. After fixation, cells were washed twice with PBS, followed by blocking with 10% normal goat serum in PBS for 1 hour at room temperature.

Cells were then incubated overnight at 4 °C in blocking buffer, either without primary antibody (no primary control) or with cadherin-11 (extracellular) rabbit polyclonal antibody (1:200, DF3523, Affinity Biosciences). After three washes with PBS, cells were incubated for 1 hour at room temperature in blocking buffer containing 1 μg/ml Hoechst 33342, BioTracker NIR680 (1:2000, Merck), and goat anti-rabbit Alexa Fluor 488 secondary antibody (1:500, Invitrogen).

After four additional PBS washes, cells were imaged in PBS using a Leica Stellaris 5 confocal microscope equipped with a 40× oil immersion objective. Tile-scanned images were stitched using Las X microscopy software (Leica) to generate high-resolution panoramic images.

### BioTracker staining

BioTracker NIR680 was diluted 1:2000 in unsupplemented DMEM/F-12. T25 flasks containing FLS were washed once with HBSS (with calcium and magnesium; Gibco), then incubated with the diluted dye for 30 minutes at 37 °C. After incubation, the flasks were washed three times with FLS media, with each wash lasting 10 minutes at 37°C. FLS were then dissociated using Versene and plated onto 6-or 24-well plates.

### FLS transfection

For transfection in 6-well plates, the medium was removed and replaced with 1 ml of Opti-MEM I (Gibco). In a microcentrifuge tube, 500 ng of saRNA was diluted in 200 μl of Opti-MEM I. In a separate tube, 3 μl of Lipofectamine MessengerMAX (Invitrogen) was diluted in 100 μl of Opti-MEM I. After gentle mixing, the solutions were combined and incubated at room temperature for 5 minutes, then added dropwise to the cells. The cells were incubated with the complexes for 2 hours at 37 °C, after which the medium was removed and replaced with fresh FLS media. For transfection in 24-well plates, all volumes and amounts of saRNA were reduced fivefold.

### Microplate imaging

Black glass-bottom 6-well plates (P06-1.5H-N, Cellvis) or 24-well plates (662892, Greiner Bio-one) were used for microplate imaging. Prior to imaging, the FLS media was replaced with Live Cell Imaging Solution (Invitrogen). Imaging was performed using an Odyssey M laser scanner (LI-COR), using LI-COR acquisition software, a plate offset of +1.45 mm, and 100 μm resolution.

EGFP, mScarlet3, and BioTracker NIR680 were imaged using the 488, 520, and 700 channels, respectively. Spectral cross-excitation between EGFP and mScarlet3 in the 488 and 520 channels was corrected through linear unmixing analysis, as described below. Fluorescence intensity was quantified in ImageJ by measuring the integrated density within equal-area regions of interest for each well.

Fluorescence values (EGFP, mScarlet3, and BioTracker) were normalized to the day 0 BioTracker signal measured before transfection to account for variations in starting cell number.

### Linear unmixing analysis

saRNA constructs encoding individual fluorescent proteins (moxBFP, EGFP, or mScarlet3) were designed to assess cross-excitation among the fluorescent proteins used in this study. tSA201 cells (96121229, ECACC) were seeded in black glass-bottom 6-well plates coated with poly-D-lysine (Sigma) and transfected with the respective saRNA constructs. Cells were imaged the following day using an Odyssey M laser scanner, with fluorescence captured through the 488 and 520 channels.

Expression of moxBFP was undetectable in both the 488 and 520 channels, and therefore it was excluded from further analysis. Bleed-through of EGFP into the 520 channel and mScarlet3 into the 488 channel was quantified using ImageJ, with cross-excitation determined to be 11.32% and 0.94%, respectively. The emission signals were assumed to be linearly proportional to the sum of the intensities of each fluorophore, and the unmixed EGFP and mScarlet3 signals were calculated using Python.

### Annexin V assay

FLS were stained with BioTracker NIR680 and plated on black glass-bottom 24-well plates (Sensoplate, Greiner Bio-One). Cells were transfected with saRNA or treated with 0.5 μM staurosporine (Cayman Chemical) as a positive control.

Annexin V-CF800 conjugate (Biotium) was diluted to 250 ng/ml in Live Cell Imaging Solution (Invitrogen). On each day of the assay, wells were washed with Live Cell Imaging Solution and replaced with diluted Annexin V-CF800 solution. Plates were incubated at 37 °C for 15 minutes. Following incubation, plates were washed three times with Live Cell Imaging Solution before imaging the 700 and 800 channels with an Odyssey M imager set to +1.45 mm image offset and 100 μm resolution.

Image analysis was performed using ImageJ. To correct for unidirectional spectral bleed-through of BioTracker NIR680 into the 800-channel, subtractive compensation was applied by dividing the 700-channel image by a factor of 800 and then subtracting it from the 800-channel image. Due to the presence of a small number of high-intensity speckles in the 800-channel image, area rather than fluorescence intensity was quantified. The display range of the 800-channel image was set between 0.25 and 2.5, and the image was thresholded using the method of Li.^136^ The area of thresholded pixels within each well was then quantified and adjusted based on the BioTracker signal before transfection to account for variations in cell number. Data were then normalized to the average of the mock transfection condition.

### Calcein AM staining

Calcein AM (Invitrogen) staining was performed according to the manufacturer’s protocol. Briefly, cells were washed with HBSS and incubated with 2 μM calcein AM (diluted in HBSS) at 37 °C for 30 minutes. Following three washes, cells were imaged in Live Cell Imaging Solution, and the 488-channel image was captured using an Odyssey M imager.

Although calcein AM and EGFP have overlapping spectra, calcein AM fluorescence was typically much higher than EGFP, making EGFP’s contribution to the measured Calcein AM signal negligible in most cases. However, when the E3 construct was used, EGFP expression was sufficient to interfere with the signal. To address this, in experiments that included use of the E3 construct, a 488-channel image was captured prior to calcein AM application and subtracted from the post-application image to accurately measure calcein AM fluorescence.

Fluorescence intensity was quantified using ImageJ. In experiments where BioTracker was used, fluorescence values were corrected by the pre-transfection BioTracker signal to account for differences in starting cell number.

### In-cell western assay

Cells were plated on black, glass-bottom 24-well plates. Alongside mock and saRNA transfections, one well was reserved for background subtraction, which received no treatment. Unless specified otherwise, experiments were conducted 2 days post-transfection. Cells were fixed with 4% paraformaldehyde for 10 minutes at room temperature, followed by two washes with tris-buffered saline (TBS). Permeabilization was then performed using 0.1% Triton X-100 in TBS for 10 minutes, followed by two additional TBS washes. After permeabilization, cells were blocked with Intercept TBS Blocking Buffer (LI-COR) for 1 hour at room temperature with gentle agitation.

Primary antibodies were diluted in Intercept Blocking Buffer and incubated with the cells overnight at 4 °C. The background subtraction well was incubated with blocking buffer alone, without primary antibodies.

The following primary antibodies were used: Phospho-eIF2α (Ser51) (D9G8) XP rabbit monoclonal (1:200, #3398, Cell Signaling Technology), eIF2α (L57A5) mouse monoclonal (1:200, #2103, Cell Signaling Technology), PKR rabbit polyclonal (1:500, #18244-1-AP, Proteintech), Phospho-eIF4E (S209) rabbit monoclonal (1:200, #ab76256, Abcam), eIF4E mouse monoclonal (5D11) (1:200, #MA1-089, Invitrogen), and fibroblast activation protein (73.3) mouse monoclonal (20 μg/ml, #BE0374, InVivoMAb). Except for the fibroblast activation protein antibody, which was conjugated to DyLight 800 using the manufacturer’s DyLight Antibody Labeling Kit (#53062, Thermo Scientific), all primary antibodies were unconjugated.

After washing the wells four times with TBS supplemented with 0.1% Tween 20 (TBST), cells were incubated for 1 hour at room temperature with the appropriate secondary antibodies or normalization stain, with gentle agitation. The secondary antibodies used were goat anti-mouse IRDye 800CW (1:800), donkey anti-rabbit IRDye 800CW (1:800), donkey anti-mouse IRDye 680RD (1:800), and CellTag 700 (1:500), all from LI-COR. The background subtraction well received secondary antibody but no CellTag. Following four additional washes with TBST, the plate was inverted and gently tapped on absorbent paper to remove excess liquid. The plate was then imaged using the Odyssey M imager with LI-COR Acquisition software using a plate offset of +1.45 and 100 μm resolution. Signal quantification was carried out using Empiria Studio software (LI-COR).

### SUrface SEnsing of Translation (SUnSET) assay

FLS were plated in black, glass-bottom 24-well plates, with one well reserved for background subtraction (no treatment). Two days post-transfection, cells were treated with 1 μM puromycin (Sigma) for 30 minutes at 37 °C. The background control well did not receive puromycin. Following incubation, cells were washed twice with HBSS and fixed in 5% neutral-buffered formalin (in PBS) for 20 minutes at room temperature. Cells were then washed three times with PBS, permeabilized with 0.2% Triton X-100 for 10 minutes, and blocked with Intercept Blocking Buffer for 1 hour at room temperature.

Anti-puromycin antibody (3RH11, Absolute Antibody) was diluted 1:250 in blocking buffer and incubated with cells overnight at 4 °C. The following day, cells were washed four times with TBST, then incubated with goat anti-mouse IRDye 800CW (1:800) and CellTag 700 (1:500) diluted in blocking buffer. The background subtraction well did not receive CellTag. After four additional washes with TBST, plates were inverted, tapped dry on absorbent paper, and imaged using the Odyssey M imager. Signal quantification was performed using Empiria Studio software.

### rRNA integrity analysis

FLS were plated in 6-well plates and either transfected with saRNA or subjected to mock transfection. After 26–30 hours (allowing for saRNA replication but before substantial cell loss), cells were harvested by scraping in 350 μl RLT buffer (QIAGEN) supplemented with 10 μl/ml β-mercaptoethanol (Sigma). Cell lysates were homogenized using QIAshredder columns (QIAGEN), and total RNA was extracted using the RNeasy Mini Kit (QIAGEN) according to the manufacturer’s instructions. RNA quantity and quality were initially assessed using a N60 nanophotometer (Implen). Samples were aliquoted and stored at-80 °C until further analysis. For detailed quality assessment, samples were transported on dry ice and analyzed using the RNA Pico Kit on the 2100 Bioanalyzer system (Agilent), performed by Cambridge Genomic Services. RNA Integrity Number (RIN) values were determined for each sample.

### Bead-based immunoassay

FLS were stained with BioTracker and seeded onto 6-well plates. After transfection, the saRNA-lipofectamine complexes were replaced with 2 ml of fresh FLS media. Two days later, supernatants were collected and stored at –80 °C until analysis. On the day of analysis, the supernatant was thawed on ice, and levels of 13 cytokines (IFN-γ, CXCL1, TNF-α, MCP-1, IL-12p70, CCL5, IL-1β, CXCL10, GM-CSF, IL-10, IFN-β, IFN-α, and IL-6) were measured using the LEGENDplex Mouse Anti-Virus Response Panel (740621, BioLegend) following the manufacturer’s protocol. Samples were run in duplicate. A custom 3D-printed PETG vacuum manifold (designed with Fusion 360) was used for vacuum filtration, and beads were analyzed using a CytoFLEX LX flow cytometer (Beckman Coulter). Data analysis was performed with the LEGENDplex Qognit Data Analysis Suite (BioLegend). To account for differences in cell number between wells, cytokine levels were normalized to the BioTracker signal before transfection.

### ML336 experiments

ML336 (Cayman Chemical) was dissolved in DMSO (Sigma) to a stock concentration of 10 mM and stored in aliquots at –20 °C. ML336 was added to cell cultures at the indicated final concentrations. Vehicle controls received either 0.01% or 0.1% DMSO, corresponding to the maximum DMSO concentrations used in the respective ML336-treated conditions.

### RT-qPCR experiments

FLS were seeded in 96-well plates, and transfections were performed using a 30-fold downscaling of the volumes used in 6-well formats. At 2 days post-transfection, cDNA was synthesized directly from cell lysates using the TaqMan Fast Advanced Cells-to-CT Kit (Invitrogen), according to the manufacturer’s protocol, and stored at –80 °C until analysis. For each biological replicate, a single well that received no treatment and was processed without reverse transcriptase (RT) served as a shared no-RT control.

For each treatment condition, a reaction mix containing cDNA, RT-PCR grade water (Invitrogen), and TaqMan™ Fast Advanced Master Mix (Applied Biosystems) was prepared and dispensed into the corresponding wells of custom 96-well 0.1 ml TaqMan array plates (ThermoFisher). All conditions for a given biological replicate were run on the same plate to minimize inter-plate variability. Plates were run in duplicate, and each plate included no-template and no-RT controls, along with cDNA from mock-transfected cells, and cells transfected with native saRNA, E3, or srIκBα-Smad7-SOCS1 constructs.

Each array included the following TaqMan Gene Expression Assays: *18S* (Hs99999901_s1), *Adar* (Mm00508001_m1), *Isg20* (Mm00469585_m1), *Rigi* (Mm01216853_m1), *Ifih1* (Mm00459183_m1), *Oas1e* (Mm00653062_m1), *Tlr3* (Mm01207404_m1), *Eif2ak2* (Mm01235643_m1), *ZC3hav1* (Mm00512227_m1), *Rnasel* (Mm00712008_m1), *EGFP* (Mr04329676_mr), *Ifnb1* (Mm00439552_s1), *Ifna2* (Mm00833961_s1), *Ifng* (Mm01168134_m1), *Tnf* (Mm00443258_m1), and *Il6* (Mm00446190_m1).

Reactions were run for 40 cycles on a StepOnePlus Real-Time PCR System (Applied Biosystems). A consistent ΔRn threshold was applied across all plates and replicates. ΔCT values were calculated relative to *18S*. Transcripts not detected in all samples were excluded from analysis, with the exception of EGFP. These excluded transcripts included *Oas1e*, *Ifnb1*, *Ifna2*, *Ifng*, and *Tnf*. Statistical significance was evaluated using ΔCT values, and mean –ΔΔCT values relative to the mock condition are shown as a heat map for visualization.

### Statistical Analysis

All statistical analyses were conducted using GraphPad Prism 9, with specific tests detailed in the corresponding figure legends. All statistical tests were two-sided tests.

## DATA AVAILABILITY STATEMENT

The code for the unmixing analysis is available at https://github.com/lariferg/spectral_unmixing.

The 3D printed vacuum manifold compatible with the LEGENDplex assay has been deposited in the NIH 3D print exchange (3DPX-021388).

All data needed to evaluate the conclusions in the paper are present in the paper, the Supplemental Materials, and the Figshare repository http://doi.org/10.6084/m9.figshare.27091972.

## ACKNOWLEDGMENTS

The authors gratefully acknowledge the Cambridge Advanced Imaging Centre, the flow cytometry facility from the School of the Biological Sciences, and Cambridge Genomic Services for their support & assistance in this work. The authors would like to thank Dr. Paul Miller for providing the pDx_mScarlet3 plasmid and Dr. Alex Cloake for providing the pIRES2-EGFP plasmid used in this study. T.K.L. acknowledges support from a Horizon Europe Marie Skłodowska-Curie Actions European Postdoctoral Fellowship (UKRI Guarantee) [EP/X023117/1]. A.R. and L.W.P. disclose support from AstraZeneca PhD studentships [G115018 and G113502, respectively]. L.F. discloses support from funding provided by the MRC Postdoctoral Training Scheme. E.St.J.S. acknowledges funding from the UKRI and Versus Arthritis [MR/W002426/1] as part of the ADVANTAGE visceral pain consortium through the Advanced Pain Discovery Platform (APDP) and, E.St.J.S. and L.J.G. acknowledge the Wellcome Trust [225856/Z/22/Z].

## AUTHOR CONTRIBUTIONS

Conceptualization, T.K.L. and E.St.J.S Formal analysis: T.K.L., E.St.J.S. Funding acquisition: E.St.J.S., T.K.L. Investigation: T.K.L., A.R., L.W.P., L.J.G. Methodology: T.K.L., T.A.

Resources: E.St.J.S., T.K.L. Software: L.F.

Supervision: E.St.J.S., T.K.L. Visualisation: T.K.L., E.St.J.S. Writing—original draft: T.K.L.

Writing—reviewing & editing: E.St.J.S., L.F.

## DECLARATION OF INTERESTS

Authors declare that they have no competing interests.

## DECLARATION OF GENERATIVE AI AND AI-ASSISTED TECHNOLOGIES IN THE WRITING PROCESS

During the preparation of this work, the authors used OpenAI’s ChatGPT to enhance clarity and flow of the writing. After using this tool, the authors reviewed and edited the content as needed and take full responsibility for the content of the publication.

---

**Figure 1—figure supplement 1.**
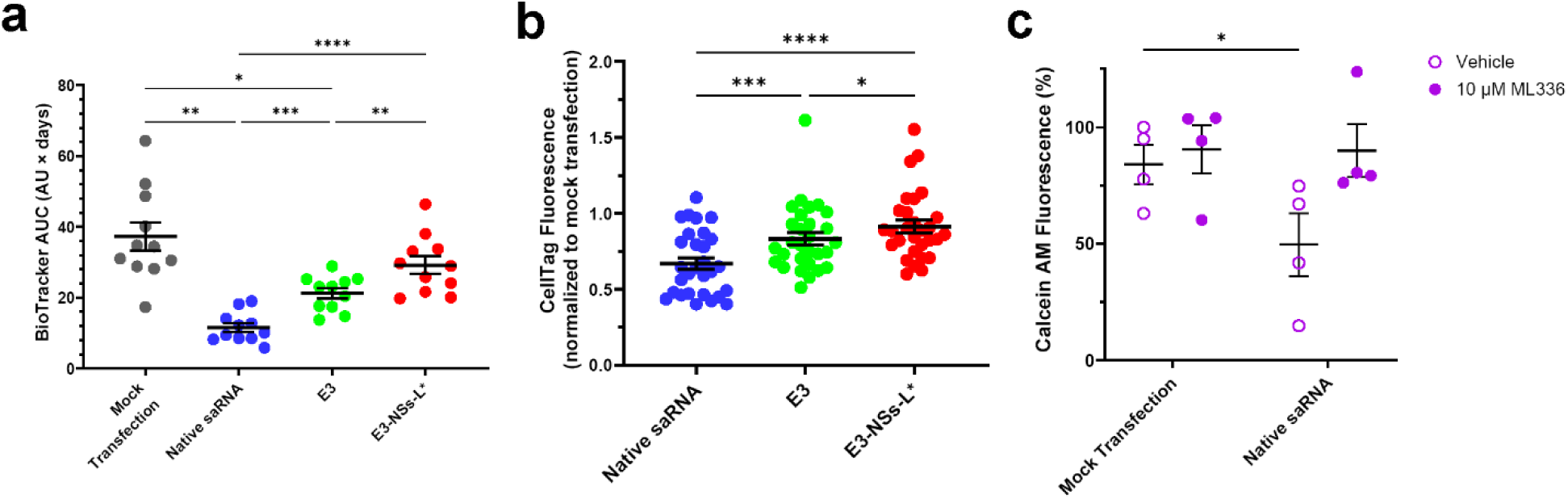
Blocking dsRNA-sensing pathways or saRNA replication inhibits saRNA-induced cell loss **a,** AUC analysis of BioTracker fluorescence data shown in Figure 1f, summarizing cumulative effects over the time course (n = 11 biological replicates). Increasing dsRNA-sensing pathway inhibition prevents saRNA-induced reductions in integrated BioTracker signal. Statistical analysis was performed using one-way RM ANOVA with Greenhouse–Geisser correction and Tukey’s multiple comparisons test to compare all groups. Mock transfection data is also presented in Supplemental Figure S5a. **b,** Quantification of mock transfection normalized CellTag signal (n = 29 biological replicates). Increasing dsRNA-sensing pathway inhibition mitigates saRNA-induced reductions in CellTag signal. Statistical analysis was performed using one-way RM ANOVA with Greenhouse–Geisser correction and Tukey’s multiple comparisons test to compare all groups. Assays were performed 2 days post-transfection. Data in this panel were pooled from in-cell western assays, incorporating data presented in Figure 3, Figure 3—figure supplement 1c, Figure 6, and additional data not shown elsewhere. **c,** Quantification of calcein AM viability dye signal before and one day after transfection with native saRNA or mock transfection, with or without the RdRp inhibitor ML336 (n = 4 biological replicates). Transfection with native saRNA reduced cell viability, an effect that was prevented by co-treatment with ML336. Statistical analysis was performed using two-way repeated-measures ANOVA with Greenhouse– Geisser correction, followed by Dunnett’s multiple comparisons test relative to the mock-transfected, vehicle-treated group. For all statistical reporting, *P < 0.05, **P < 0.01, ***P < 0.001 and ****P < 0.0001. Data are presented as mean ± SEM.

**Figure 3—figure supplement 1.**
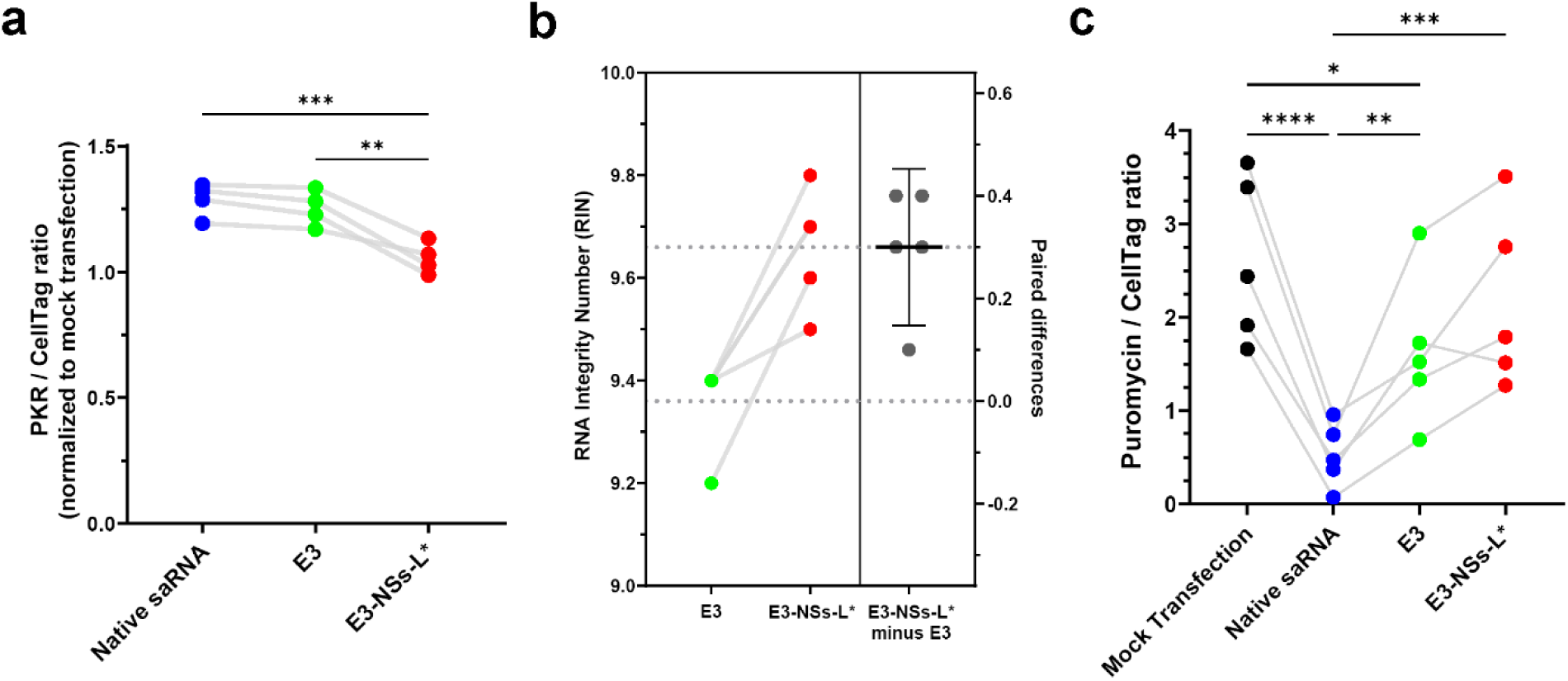
The E3-NSs-L* construct protects against saRNA-induced PKR upregulation, RNA degradation, and reduced global protein synthesis **a,** PKR levels examined day 2 post-transfection by in-cell western assay (n = 4 biological replicates). E3-NSs-L* significantly reduced PKR levels compared to both native saRNA and E3 transfection. Data are presented as fold-change relative to mock-transfected cells. Statistical significance was determined by one-way RM ANOVA with Tukey’s multiple comparisons test. **b,** rRNA integrity of FLS transfected with E3-NSs-L* is significantly higher than that of E3-transfected FLS (n = 5 biological replicates). rRNA integrity was assessed using the RNA Integrity Number (RIN) algorithm, which ranges from 1 to 10, with 10 indicating fully intact rRNA. Total RNA was extracted from FLS 1 day post-transfection. Data are shown as a Gardner-Altman comparison plot, with the right panel illustrating the mean effect size ± 95% CI. Statistical significance was assessed using a paired t-test (P = 0.0054). Dotted lines indicate group means. **c,** Translation rates were assessed two days post-transfection by measuring puromycin incorporation by in-cell western assay (n = 5 biological replicates). Transfection with native saRNA or the E3 construct significantly reduced the rate of protein synthesis, whereas the E3-NSs-L* construct had no effect. Statistical significance was determined by one-way repeated-measures ANOVA with Tukey’s multiple comparisons test. Statistical significance is indicated as follows: *P < 0.05, **P < 0.01, ***P < 0.001, and ****P < 0.0001.

## SUPPLEMENTAL FIGURES

**Supplemental Figure S1.**
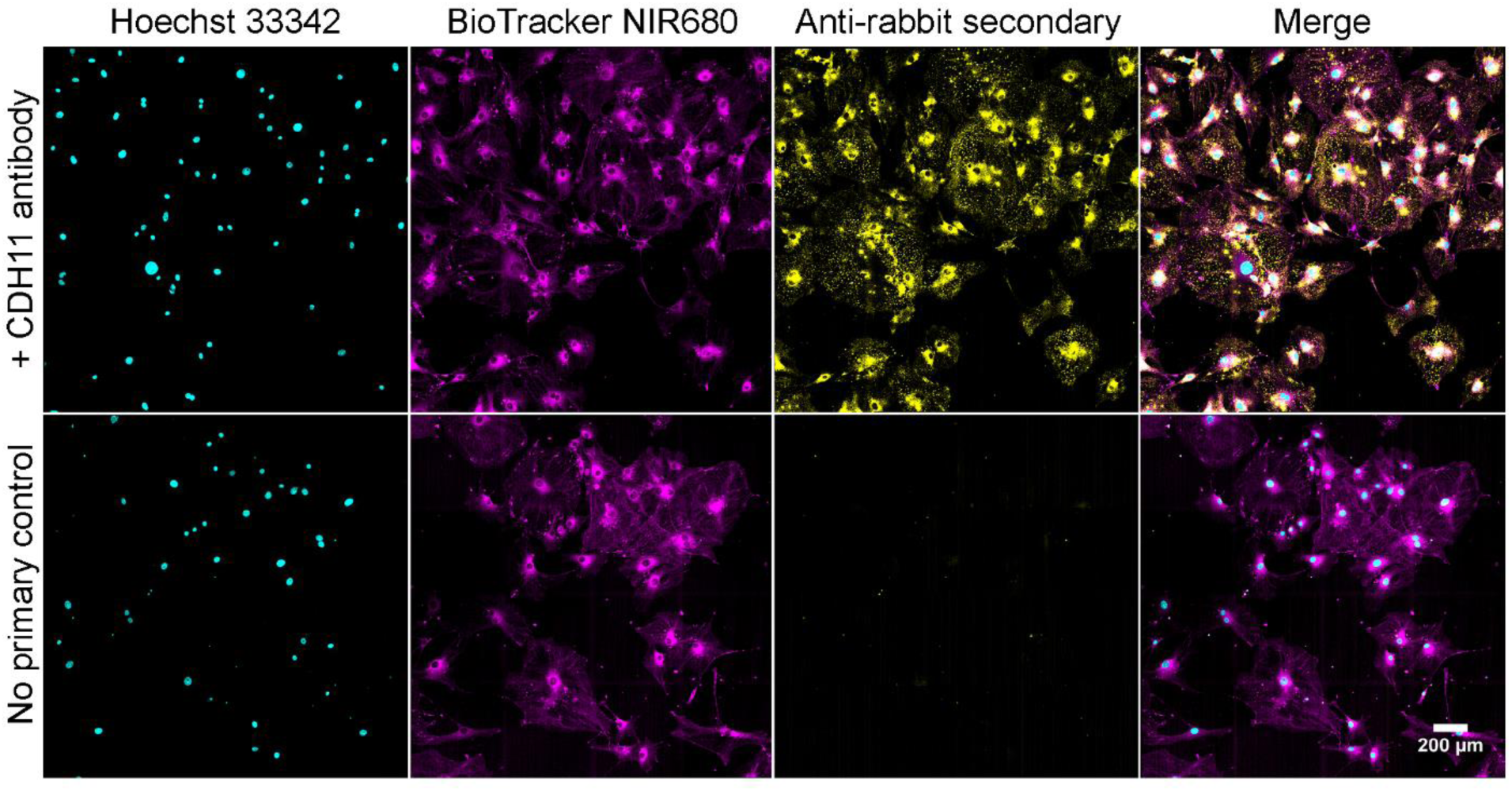
Cadherin-11, a marker of fibroblast-like synoviocytes (FLS), stains cells isolated from mouse knee explants Representative microscopy images of FLS isolated from mouse knee explants, stained with cadherin-11 (CDH11), an FLS marker. From left to right, columns show nuclear staining (Hoechst 33342, cyan), lipophilic membrane dye labelling (Biotracker NIR680, magenta), and Alexa 488 anti-rabbit secondary immunostaining (yellow), followed by a merged image. The top row shows cells treated with rabbit anti-CDH11 primary antibody, confirming CDH11 expression in isolated cells. The bottom row presents a no primary antibody control, showing no signal in the 488 channel. Scale bar = 200 μm.

**Supplemental Figure S2.**
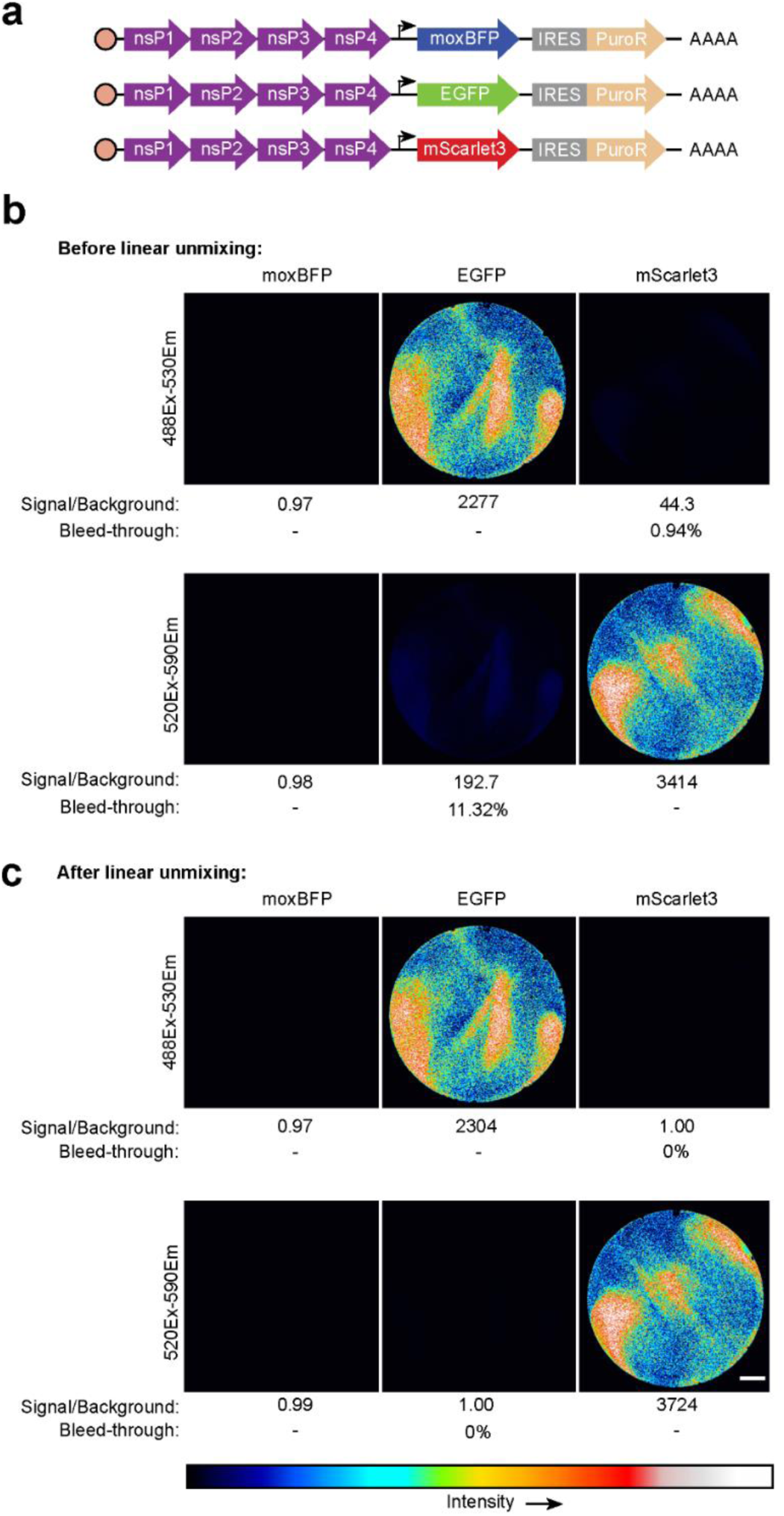
Linear unmixing corrects spectral overlap of EGFP and mScarlet3 fluorescent signals imaged using the Odyssey M laser scanner Schematic of the saRNA constructs used for linear unmixing, each expressing a different fluorescent protein: moxBFP, EGFP, and mScarlet3. These constructs were transfected into tSA201 cells. **b,** Representative imaging results showing fluorescence in the 488 and 520 channels, 1 day after transfection. moxBFP-transfected cells were not detected in either channel. EGFP was primarily detected in the 488 channel, with 11.32% bleedthrough into the 520 channel. mScarlet3 was primary detected in the 520 channel, with 0.94% bleedthrough into the 488 channel. Background signal was determined from mock transfected cells. **c,** Corrected images using linear unmixing to negate spectral bleedthrough from mScarlet3 and EGFP signals. Scale bars = 5 mm.

**Supplemental Figure S3.**
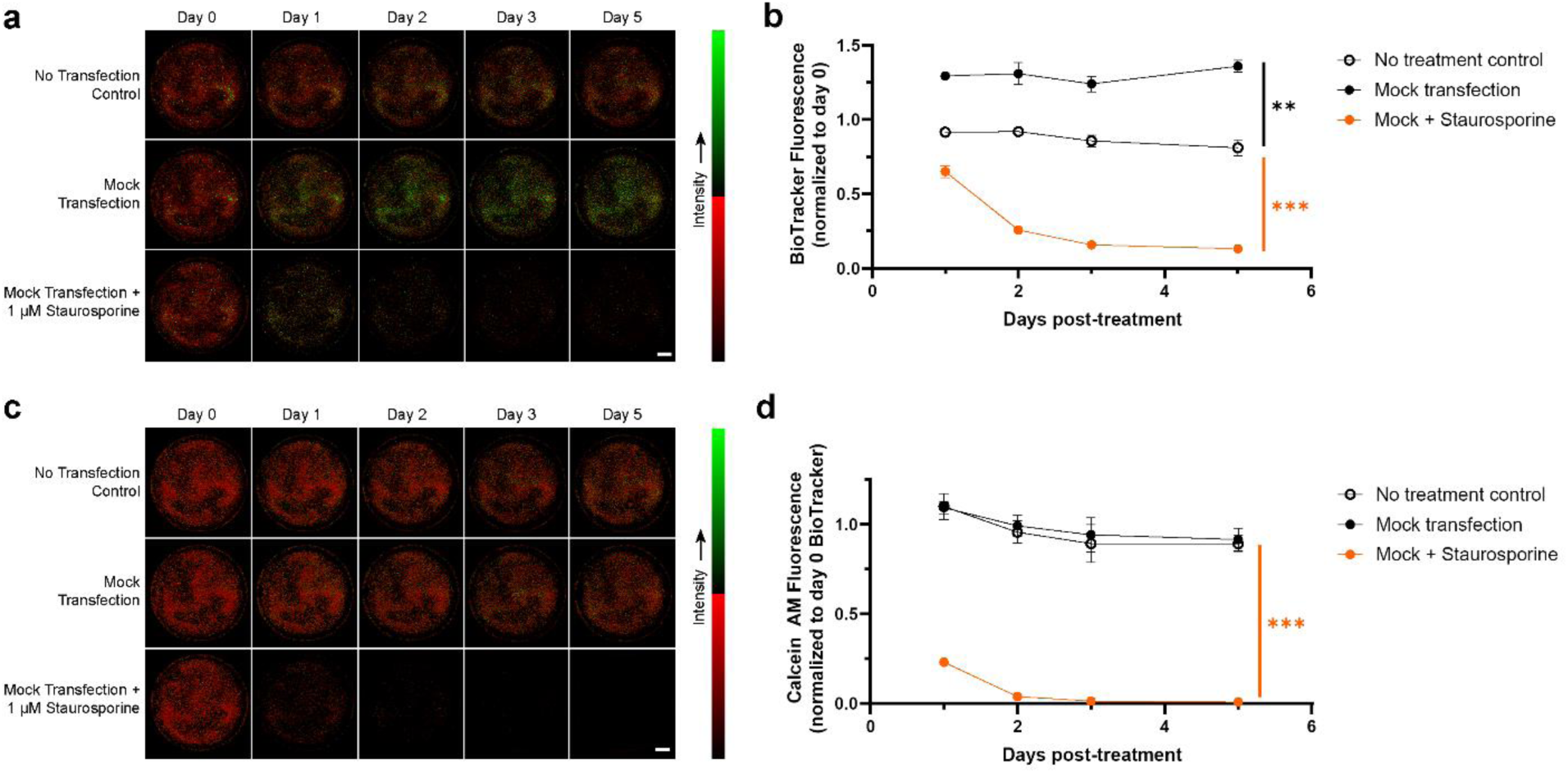
BioTracker fluorescence indicates cell number following staurosporine-induced apoptosis, despite an increase after mock transfection Validation of BioTracker fluorescence as a marker for cell number in the presence of apoptotic stimuli. **a,** Representative images of FLS stained with BioTracker under three conditions: no transfection control, mock transfection, and mock transfection with 1 μM staurosporine. BioTracker fluorescence intensity increases after mock transfection but decreases following staurosporine treatment, indicating cell loss. Scale bar = 5 mm. **b,** Quantification of BioTracker fluorescence over time for the three conditions (n = 3). Statistical significance was determined by two-way RM ANOVA with Dunnett’s multiple comparisons test comparing groups to the no treatment control. **c,** Representative images of the same FLS stained with Calcein AM. Unlike BioTracker, Calcein AM fluorescence does not increase after mock transfection but decreases after staurosporine treatment, reflecting apoptotic cell death. Scale bar = 5 mm. **d,** Quantification of BioTracker fluorescence over time for the three conditions (n = 3). Fluorescence decreases following staurosporine treatment, consistent with findings in (a, b). Statistical significance was determined by two-way RM ANOVA with Dunnett’s multiple comparisons test comparing groups to the no treatment control. For all statistical reporting, **P < 0.01 and ***P < 0.001. Data are presented as mean ± SEM.

**Supplemental Figure S4.**
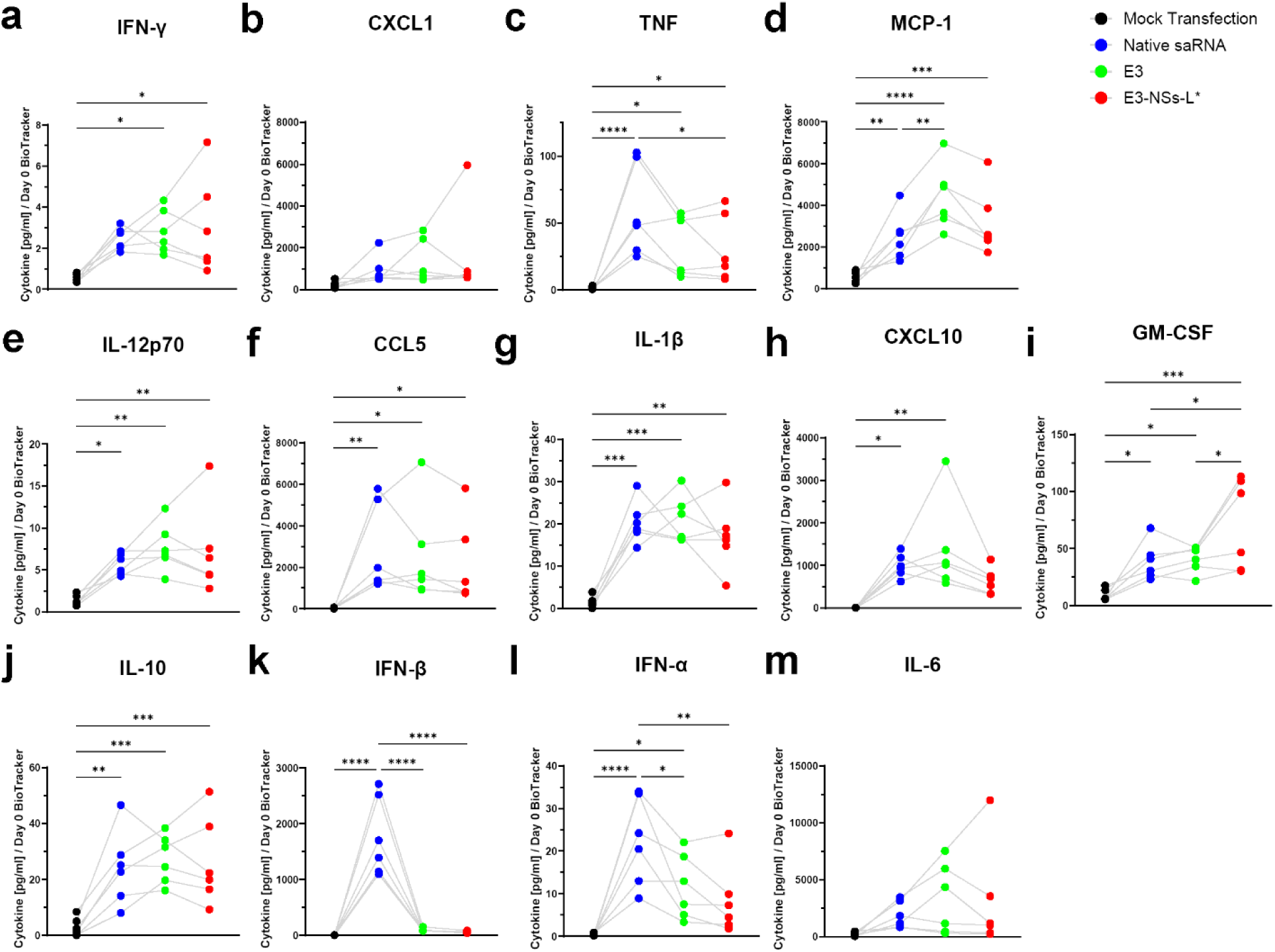
Select cytokines show variable responses to dsRNA-sensing pathway inhibition following saRNA transfection **a-m,** FLS were transfected with the saRNA constructs described in Figure 1a or mock transfected. Cell culture supernatants were collected two days post-transfection, and cytokine levels were quantified by bead-based immunoassay and normalized to cell number using the BioTracker signal measured on day 0 (n = 6 biological replicates). Native saRNA triggered robust secretion of multiple antiviral cytokines. Co-expression of viral innate immune inhibitors reduced some cytokine responses and enhanced others. Both E3 and E3-NSs-L* significantly reduced secretion of IFN-α and IFN-β, and E3-NSs-L* additionally suppressed TNF. In contrast, E3 increased MCP-1, and E3-NSs-L* increased GM-CSF levels relative to native saRNA. Statistical significance was assessed using one-way repeated-measures ANOVA for each cytokine. Multiple comparisons were controlled using the Benjamini–Krieger–Yekutieli false discovery rate (FDR) procedure (Q = 5%). All cytokines passed the discovery threshold. Treatment effects were analyzed using Tukey’s multiple comparisons test. *P < 0.05, **P < 0.01, ***P < 0.001, ****P < 0.0001. Data are presented as mean ± SEM. Mock transfection controls are shared with Supplemental Figure S5. Acronyms: IFN-γ, interferon-γ; CXCL1, C-X-C motif chemokine ligand 1; TNF, tumor necrosis factor; MCP-1, monocyte chemoattractant protein-1; IL-12p70, interleukin-12; CCL5, chemokine ligand 5; IL-1β, interleukin-1β; CXCL10, C-X-C motif chemokine ligand 10; GM-CSF, granulocyte-macrophage colony-stimulating factor; IL-10, interleukin-10; IFN-β, interferon-β; IFN-α, interferon-α; IL-6, interleukin-6.

**Supplemental Figure S5.**
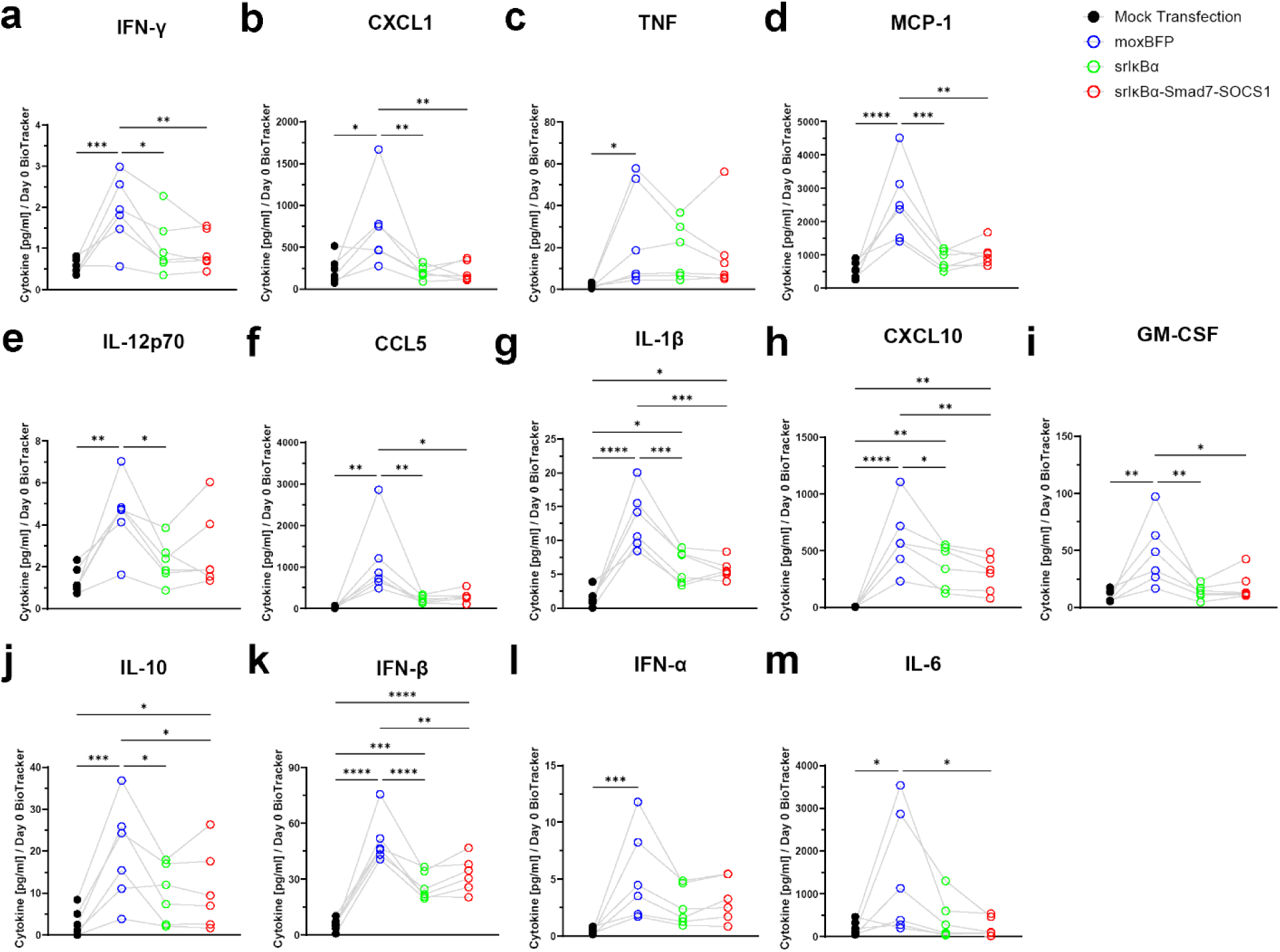
Inhibition of inflammatory signalling pathways broadly reduces saRNA-induced secretion of anti-viral cytokines **a-m,** FLS were transfected with the saRNA constructs described in Figure 4b or mock transfected. Cell culture supernatants were collected two days post-transfection, and cytokine levels were quantified by bead-based immunoassay and normalized to cell number using the BioTracker signal measured on day 0 (n = 6 biological replicates). The moxBFP construct triggered secretion of antiviral cytokines, which were broadly reduced by expression of srIκBα or srIκBα-Smad7-SOCS1. Statistical significance was assessed using one-way repeated-measures ANOVA for each cytokine. Multiple comparisons were controlled using the Benjamini–Krieger–Yekutieli false discovery rate (FDR) procedure (Q = 5%). All cytokines passed the discovery threshold. Treatment effects were analyzed using Tukey’s multiple comparisons test. *P < 0.05, **P < 0.01, ***P < 0.001, ****P < 0.0001. Data are presented as mean ± SEM. Mock transfection controls are shared with Supplemental Figure S4. Acronyms: IFN-γ, interferon-γ; CXCL1, C-X-C motif chemokine ligand 1; TNF, tumor necrosis factor; MCP-1, monocyte chemoattractant protein-1; IL-12p70, interleukin-12; CCL5, chemokine ligand 5; IL-1β, interleukin-1β; CXCL10, C-X-C motif chemokine ligand 10; GM-CSF, granulocyte-macrophage colony-stimulating factor; IL-10, interleukin-10; IFN-β, interferon-β; IFN-α, interferon-α; IL-6, interleukin-6.

**Supplemental Figure S6.**
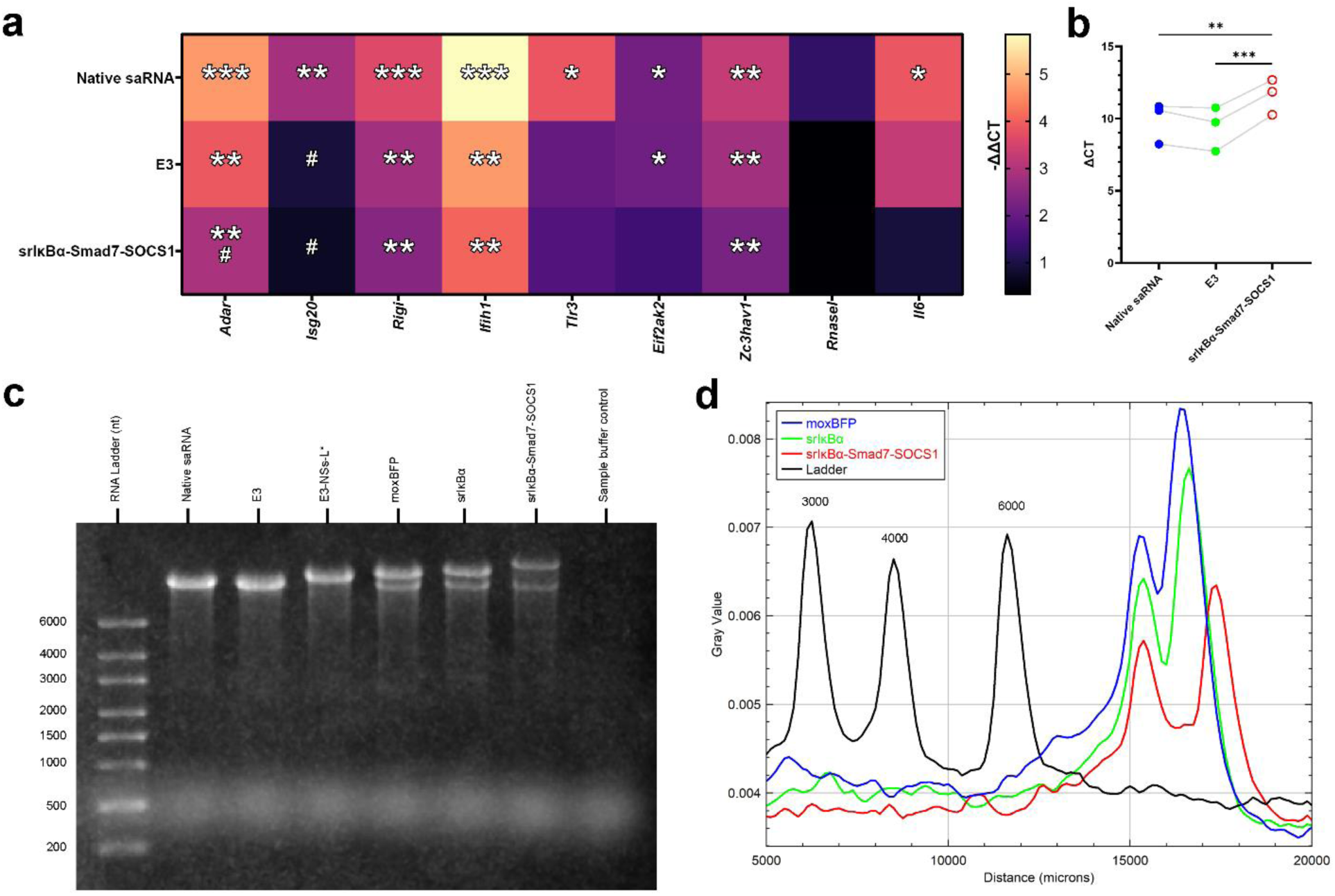
Reduced antiviral gene expression and replicon activity observed with co-expression of inflammatory signalling inhibitors **a,** Antiviral and proinflammatory transcripts were quantified by qPCR 2 days post-transfection (n = 3 biological replicates). Statistical comparisons were performed on ΔCT values normalized to *18S* rRNA using one-way ANOVA with multiple comparisons controlled using the two-stage linear step-up procedure of Benjamini, Krieger, and Yekutieli (FDR = 5%). All investigated transcripts passed significance thresholds for discovery. Treatment effects were analyzed using Holm-Šídák’s multiple comparisons test. *P < 0.05, **P < 0.01, ***P < 0.001 vs. mock transfection; ^#^P < 0.05 vs. native saRNA. Mean-ΔΔCT values normalized to mock transfection are shown on the heatmap (larger values indicate higher expression). **b,** EGFP transcript levels were quantified by qPCR at 2 days post-transfection (n = 3 biological replicates). Expression was normalized to *18S* rRNA, and data are shown as ΔCT values. FLS transfected with the srIκBα-Smad7-SOCS1 construct had higher ΔCT values (indicating lower EGFP transcript levels) compared to those transfected with native saRNA or the E3 construct. Statistical comparisons were performed on ΔCT values using one-way ANOVA with Tukey’s multiple comparisons test. **c,** In vitro transcribed saRNA constructs were resolved by denaturing gel electrophoresis. Native saRNA (11,181 nt), E3 (11,030 nt), and E3-NSs-L* (12,562 nt) migrated as single bands corresponding to their expected sizes. In contrast, the moxBFP (13,282 nt), srIκBα (13,614 nt), and srIκBα-Smad7-SOCS1 (15,664 nt) constructs each exhibited two bands: one at the expected size and a smaller lower intensity band of consistent size across all three constructs, suggestive of a common truncated transcript. **d,** Band intensity plots of moxBFP, srIκBα and srIκBα-Smad7-SOCS1 derived from the gel shown in panel (c).

**Supplemental Figure S7.**
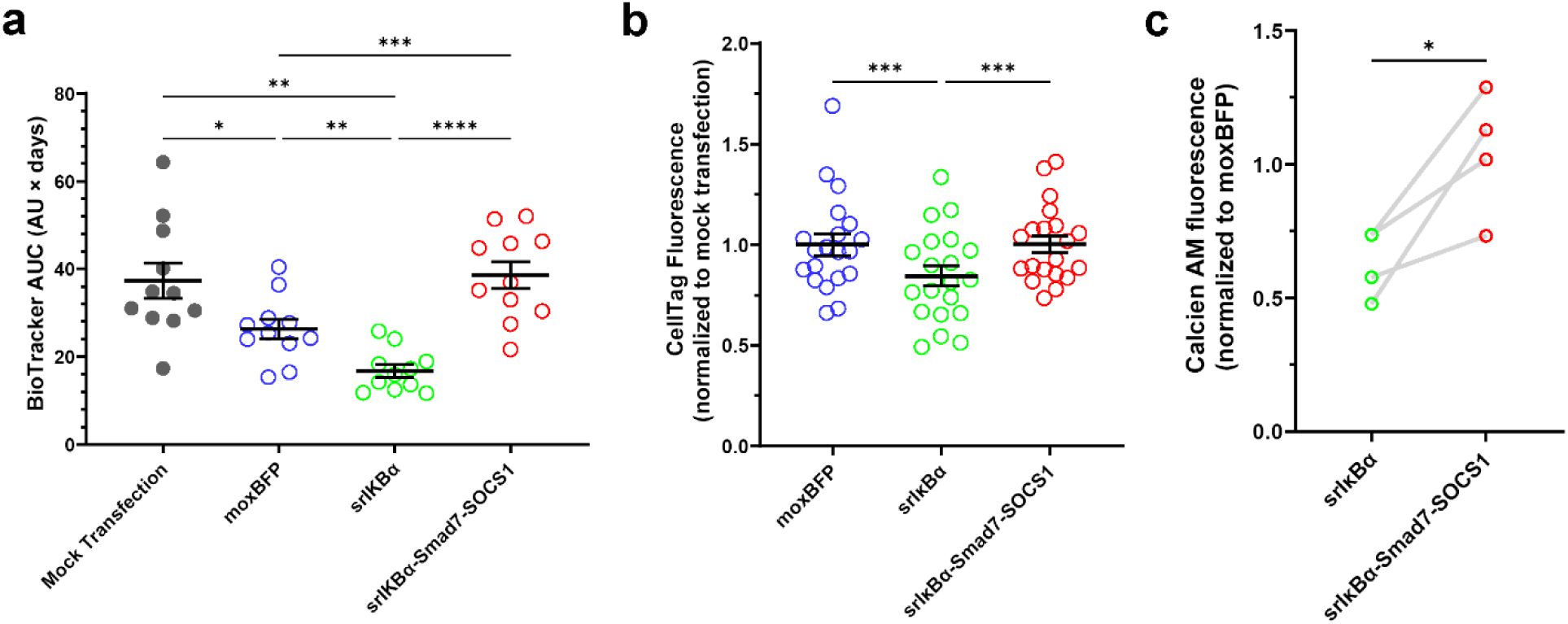
srIκBα reduces cell number and viability, while srIκBα-Smad7-SOCS1 preserves both **a,** AUC analysis of BioTracker fluorescence data shown in Figure 5c, summarizing cumulative effects over the time course (n = 11). srIκBα induces reductions in integrated BioTracker signal, which is prevented by srIκBα-Smad7-SOCS1. Statistical analysis was performed using one-way RM ANOVA with Greenhouse–Geisser correction and Tukey’s multiple comparisons test to compare all groups. Mock transfection data is also presented in Figure 1—figure supplement 1a. **b**, Quantification of mock transfection normalized CellTag signal (n = 20). The srIκBα construct reduces cell number, whereas this reduction is prevented by the srIκBα-Smad7-SOCS1 construct. Data were normalized to mock transfection. Statistical significance was determined by one-way RM ANOVA with Greenhouse–Geisser correction, and Tukey’s multiple comparisons test. Data in this panel were pooled from in-cell western assays performed on day 2 post-transfection, incorporating data presented in Figure 3 and Figure 6 as well as additional data not shown elsewhere. ***P < 0.001. **c,** Calcein AM staining performed 3 days post-transfection (n = 4). The srIκBα-Smad7-SOCS1 construct protects against reductions in cell viability induced by srIκBα. Data were normalized to the moxBFP construct. Statistical significance was determined by a paired t test. *P<0.05. Data are presented as mean ± SEM. saRNA constructs used are shown in Figure 4b.

## Notes

### Competing Interest Statement

The authors have declared no competing interest.

### Summary of Updates

Key updates include: - A puromycin assay to assess the impact of saRNA transfection and dsRNA-sensing pathway inhibition on protein synthesis; - A concentration-response analysis for ML336; - qPCR experiments examining saRNA replication efficiency and inflammatory transcript induction; - Denaturing gel electrophoresis of in vitro transcription products; - Updates to the title, graphical abstract, results and discussion sections to better reflect and communicate the findings.

## REFERENCES

1. Frolov, I., Hoffman, T.A., Prágai, B.M., Dryga, S.A., Huang, H.V., Schlesinger, S., and Rice, C.M. (1996). Alphavirus-based expression vectors: strategies and applications. Proc Natl Acad Sci U S A 93, 11371–11377.

2. Bloom, K., van den Berg, F. and Arbuthnot, P. (2021). Self-amplifying RNA vaccines for infectious diseases. Gene Ther 28, 117–129. 10.1038/s41434-020-00204-y.

3. Pietilä, M.K., Hellström, K., and Ahola, T. (2017). Alphavirus polymerase and RNA replication. Virus Research 234, 44–57. 10.1016/j.virusres.2017.01.007.

4. Maruggi, G., Zhang, C., Li, J., Ulmer, J.B., and Yu, D. (2019). mRNA as a Transformative Technology for Vaccine Development to Control Infectious Diseases. Mol Ther 27, 757–772. 10.1016/j.ymthe.2019.01.020.

5. Akhrymuk, I., Frolov, I., and Frolova, E.I. (2016). Both RIG-I and MDA5 detect alphavirus replication in concentration-dependent mode. Virology 487, 230–241. 10.1016/j.virol.2015.09.023.

6. Gong, Y., Yong, D., Liu, G., Xu, J., Ding, J., and Jia, W. (2024). A Novel Self-Amplifying mRNA with Decreased Cytotoxicity and Enhanced Protein Expression by Macrodomain Mutations. Advanced Science n/a, 2402936. 10.1002/advs.202402936.

7. Gorchakov, R., Frolova, E., Williams, B.R.G., Rice, C.M., and Frolov, I. (2004). PKR-Dependent and - Independent Mechanisms Are Involved in Translational Shutoff during Sindbis Virus Infection. Journal of Virology 78, 8455–8467. 10.1128/jvi.78.16.8455-8467.2004.

8. Dominguez, F., Palchevska, O., Frolova, E.I., and Frolov, I. (2023). Alphavirus-based replicons demonstrate different interactions with host cells and can be optimized to increase protein expression. Journal of Virology 97, e01225–23. 10.1128/jvi.01225-23.

9. Vanluchene, H., Gillon, O., Peynshaert, K., De Smedt, S.C., Sanders, N., Raemdonck, K., and Remaut, K. (2024). Less is more: Self-amplifying mRNA becomes self-killing upon dose escalation in immune-competent retinal cells. European Journal of Pharmaceutics and Biopharmaceutics 196, 114204. 10.1016/j.ejpb.2024.114204.

10. Frolov, I., and Schlesinger, S. (1994). Comparison of the effects of Sindbis virus and Sindbis virus replicons on host cell protein synthesis and cytopathogenicity in BHK cells. J Virol 68, 1721–1727. 10.1128/jvi.68.3.1721-1727.1994.

11. Terenzi, F., deVeer, M.J., Ying, H., Restifo, N.P., Williams, B.R., and Silverman, R.H. (1999). The antiviral enzymes PKR and RNase L suppress gene expression from viral and non-viral based vectors. Nucleic Acids Res 27, 4369–4375. 10.1093/nar/27.22.4369.

12. Li, Y., Su, Z., Zhao, W., Zhang, X., Momin, N., Zhang, C., Wittrup, K.D., Dong, Y., Irvine, D.J., and Weiss, R. (2020). Multifunctional oncolytic nanoparticles deliver self-replicating IL-12 RNA to eliminate established tumors and prime systemic immunity. Nat Cancer 1, 882–893. 10.1038/s43018-020-0095-6.

13. Venticinque, L., and Meruelo, D. (2010). Sindbis viral vector induced apoptosis requires translational inhibition and signaling through Mcl-1 and Bak. Mol Cancer 9, 37. 10.1186/1476-4598-9-37.

14. Mastrangelo, A.J., Hardwick, J.M., Bex, F., and Betenbaugh, M.J. (2000). Part I. Bcl-2 and Bcl-x(L) limit apoptosis upon infection with alphavirus vectors. Biotechnol Bioeng 67, 544–554.

15. Frolov, I., Agapov, E., Thomas A Hoffman, J., Prágai, B.M., Lippa, M., Schlesinger, S., and Rice, C.M. (1999). Selection of RNA Replicons Capable of Persistent Noncytopathic Replication in Mammalian Cells. Journal of Virology 73, 3854. 10.1128/jvi.73.5.3854-3865.1999.

16. Lundstrom, K., Abenavoli, A., Malgaroli, A., and Ehrengruber, M.U. (2003). Novel semliki forest virus vectors with reduced cytotoxicity and temperature sensitivity for long-term enhancement of transgene expression. Molecular Therapy 7, 202–209. 10.1016/S1525-0016(02)00056-4.

17. Pepini, T., Pulichino, A.-M., Carsillo, T., Carlson, A.L., Sari-Sarraf, F., Ramsauer, K., Debasitis, J.C., Maruggi, G., Otten, G.R., Geall, A.J., et al. (2017). Induction of an IFN-Mediated Antiviral Response by a Self-Amplifying RNA Vaccine: Implications for Vaccine Design. The Journal of Immunology 198, 4012–4024. 10.4049/jimmunol.1601877.

18. Blakney, A.K., McKay, P.F., Bouton, C.R., Hu, K., Samnuan, K., and Shattock, R.J. (2021). Innate Inhibiting Proteins Enhance Expression and Immunogenicity of Self-Amplifying RNA. Mol Ther 29, 1174–1185. 10.1016/j.ymthe.2020.11.011.

19. McGee, J.E., Kirsch, J.R., Kenney, D., Cerbo, F., Chavez, E.C., Shih, T.-Y., Douam, F., Wong, W.W., and Grinstaff, M.W. (2024). Complete substitution with modified nucleotides in self-amplifying RNA suppresses the interferon response and increases potency. Nat Biotechnol, 1–7. 10.1038/s41587-024-02306-z.

20. Zhong, Z., McCafferty, S., Opsomer, L., Wang, H., Huysmans, H., De Temmerman, J., Lienenklaus, S., Portela Catani, J.P., Combes, F., and Sanders, N.N. (2021). Corticosteroids and cellulose purification improve, respectively, the in vivo translation and vaccination efficacy of sa-mRNAs. Molecular Therapy 29, 1370–1381. 10.1016/j.ymthe.2021.01.023.

21. Beissert, T., Koste, L., Perkovic, M., Walzer, K.C., Erbar, S., Selmi, A., Diken, M., Kreiter, S., Türeci, Ö., and Sahin, U. (2017). Improvement of In Vivo Expression of Genes Delivered by Self-Amplifying RNA Using Vaccinia Virus Immune Evasion Proteins. Hum Gene Ther 28, 1138–1146. 10.1089/hum.2017.121.

22. Yoshioka, N., Gros, E., Li, H.-R., Kumar, S., Deacon, D.C., Maron, C., Muotri, A.R., Chi, N.C., Fu, X.-D., Yu, B.D., et al. (2013). Efficient Generation of Human iPS Cells by a Synthetic Self-Replicative RNA. Cell Stem Cell 13, 10.1016/j.stem.2013.06.001.

23. Xue, Y., Zhang, Y., Zhong, Y., Du, S., Hou, X., Li, W., Li, H., Wang, S., Wang, C., Yan, J., et al. (2024). LNP-RNA-engineered adipose stem cells for accelerated diabetic wound healing. Nat Commun 15, 739. 10.1038/s41467-024-45094-5.

24. Li, Y., Teague, B., Zhang, Y., Su, Z., Porter, E., Dobosh, B., Wagner, T., Irvine, D.J., and Weiss, R. (2019). In vitro evolution of enhanced RNA replicons for immunotherapy. Sci Rep 9, 6932. 10.1038/s41598-019-43422-0.

25. Blakney, A.K., Ip, S., and Geall, A.J. (2021). An Update on Self-Amplifying mRNA Vaccine Development. Vaccines (Basel) 9, 97. 10.3390/vaccines9020097.

26. Erasmus, J.H., Archer, J., Fuerte-Stone, J., Khandhar, A.P., Voigt, E., Granger, B., Bombardi, R.G., Govero, J., Tan, Q., Durnell, L.A., et al. (2020). Intramuscular Delivery of Replicon RNA Encoding ZIKV-117 Human Monoclonal Antibody Protects against Zika Virus Infection. Molecular Therapy Methods & Clinical Development 18, 402–414. 10.1016/j.omtm.2020.06.011.

27. Kim, Y.G., Baltabekova, A.Z., Zhiyenbay, E.E., Aksambayeva, A.S., Shagyrova, Z.S., Khannanov, R., Ramanculov, E.M., and Shustov, A.V. (2017). Recombinant Vaccinia virus-coded interferon inhibitor B18R: Expression, refolding and a use in a mammalian expression system with a RNA-vector. PLOS ONE 12, e0189308. 10.1371/journal.pone.0189308.

28. Merrick, W.C. (2004). Cap-dependent and cap-independent translation in eukaryotic systems. Gene 332, 1–11. 10.1016/j.gene.2004.02.051.

29. Yang, Y., and Wang, Z. (2019). IRES-mediated cap-independent translation, a path leading to hidden proteome. Journal of Molecular Cell Biology 11, 911–919. 10.1093/jmcb/mjz091.

30. Maglaviceanu, A., Wu, B., and Kapoor, M. (2021). Fibroblast-like synoviocytes: Role in synovial fibrosis associated with osteoarthritis. Wound Repair and Regeneration 29, 642–649. 10.1111/wrr.12939.

31. Mousavi, M.J., Karami, J., Aslani, S., Tahmasebi, M.N., Vaziri, A.S., Jamshidi, A., Farhadi, E., and Mahmoudi, M. (2021). Transformation of fibroblast-like synoviocytes in rheumatoid arthritis; from a friend to foe. Autoimmun Highlights 12, 3. 10.1186/s13317-020-00145-x.

32. Honig, M.G., and Hume, R.I. (1986). Fluorescent carbocyanine dyes allow living neurons of identified origin to be studied in long-term cultures. J Cell Biol 103, 171–187. 10.1083/jcb.103.1.171.

33. Bertrand, R., Solary, E., O’Connor, P., Kohn, K.W., and Pommier, Y. (1994). Induction of a Common Pathway of Apoptosis by Staurosporine. Experimental Cell Research 211, 314–321. 10.1006/excr.1994.1093.

34. Rehwinkel, J., and Gack, M.U. (2020). RIG-I-like receptors: their regulation and roles in RNA sensing. Nat Rev Immunol 20, 537–551. 10.1038/s41577-020-0288-3.

35. Junior, A.G.D., Sampaio, N.G., and Rehwinkel, J. (2019). A Balancing Act: MDA5 in Antiviral Immunity and Autoinflammation. Trends in Microbiology 27, 75–85. 10.1016/j.tim.2018.08.007.

36. Gal-Ben-Ari, S., Barrera, I., Ehrlich, M., and Rosenblum, K. (2019). PKR: A Kinase to Remember. Front. Mol. Neurosci. 11. 10.3389/fnmol.2018.00480.

37. Silverman, R.H. (2007). Viral Encounters with 2′,5′-Oligoadenylate Synthetase and RNase L during the Interferon Antiviral Response. Journal of Virology 81, 12720–12729. 10.1128/jvi.01471-07.

38. Szczerba, M., Subramanian, S., Trainor, K., McCaughan, M., Kibler, K.V., and Jacobs, B.L. (2022). Small Hero with Great Powers: Vaccinia Virus E3 Protein and Evasion of the Type I IFN Response. Biomedicines 10, 235. 10.3390/biomedicines10020235.

39. Kalveram, B., and Ikegami, T. (2013). Toscana Virus NSs Protein Promotes Degradation of Double-Stranded RNA-Dependent Protein Kinase. J Virol 87, 3710–3718. 10.1128/JVI.02506-12.

40. Drappier, M., Jha, B.K., Stone, S., Elliott, R., Zhang, R., Vertommen, D., Weiss, S.R., Silverman, R.H., and Michiels, T. (2018). A novel mechanism of RNase L inhibition: Theiler’s virus L* protein prevents 2-5A from binding to RNase L. PLoS Pathog 14, e1006989. 10.1371/journal.ppat.1006989.

41. Liu, Z., Chen, O., Wall, J.B.J., Zheng, M., Zhou, Y., Wang, L., Ruth Vaseghi, H., Qian, L., and Liu, J. (2017). Systematic comparison of 2A peptides for cloning multi-genes in a polycistronic vector. Sci Rep 7, 2193. 10.1038/s41598-017-02460-2.

42. Baird, T.D., and Wek, R.C. (2012). Eukaryotic Initiation Factor 2 Phosphorylation and Translational Control in Metabolism. Advances in Nutrition 3, 307. 10.3945/an.112.002113.

43. Koch, A., Aguilera, L., Morisaki, T., Munsky, B., and Stasevich, T.J. (2020). Quantifying the dynamics of IRES and cap translation with single-molecule resolution in live cells. Nat Struct Mol Biol 27, 1095–1104. 10.1038/s41594-020-0504-7.

44. Baiersdörfer, M., Boros, G., Muramatsu, H., Mahiny, A., Vlatkovic, I., Sahin, U., and Karikó, K. (2019). A Facile Method for the Removal of dsRNA Contaminant from In Vitro-Transcribed mRNA. Mol Ther Nucleic Acids 15, 26–35. 10.1016/j.omtn.2019.02.018.

45. Chung, D., Schroeder, C.E., Sotsky, J., Yao, T., Roy, S., Smith, R.A., Tower, N.A., Noah, J.W., McKellip, S., Sosa, M., et al. (2010). ML336: Development of Quinazolinone-Based Inhibitors Against Venezuelan Equine Encephalitis Virus (VEEV). In Probe Reports from the NIH Molecular Libraries Program (National Center for Biotechnology Information (US)).

46. Costigan, A., Hollville, E., and Martin, S.J. (2023). Discriminating Between Apoptosis, Necrosis, Necroptosis, and Ferroptosis by Microscopy and Flow Cytometry. Current Protocols 3, e951. 10.1002/cpz1.951.

47. Liu, Q., Huang, J., Xia, J., Liang, Y., and Li, G. (2022). Tracking tools of extracellular vesicles for biomedical research. Front. Bioeng. Biotechnol. 10. 10.3389/fbioe.2022.943712.

48. Chiaradia, E., Tancini, B., Emiliani, C., Delo, F., Pellegrino, R.M., Tognoloni, A., Urbanelli, L., and Buratta, S. (2021). Extracellular Vesicles under Oxidative Stress Conditions: Biological Properties and Physiological Roles. Cells 10, 1763. 10.3390/cells10071763.

49. van Genderen, H., Kenis, H., Lux, P., Ungeth, L., Maassen, C., Deckers, N., Narula, J., Hofstra, L., and Reutelingsperger, C. (2006). In vitro measurement of cell death with the annexin A5 affinity assay. Nat Protoc 1, 363–367. 10.1038/nprot.2006.55.

50. Crowley, L.C., Marfell, B.J., Scott, A.P., and Waterhouse, N.J. (2016). Quantitation of Apoptosis and Necrosis by Annexin V Binding, Propidium Iodide Uptake, and Flow Cytometry. Cold Spring Harb Protoc 2016. 10.1101/pdb.prot087288.

51. Chan, L.L., Wilkinson, A.R., Paradis, B.D., and Lai, N. (2012). Rapid Image-based Cytometry for Comparison of Fluorescent Viability Staining Methods. J Fluoresc 22, 1301–1311. 10.1007/s10895-012-1072-y.

52. Mars, J.-C., Culjkovic-Kraljacic, B., and Borden, K.L.B. (2024). eIF4E orchestrates mRNA processing, RNA export and translation to modify specific protein production. Nucleus 15, 2360196. 10.1080/19491034.2024.2360196.

53. Burke, J.M., Moon, S.L., Matheny, T., and Parker, R. (2019). RNase L Reprograms Translation by Widespread mRNA Turnover Escaped by Antiviral mRNAs. Molecular Cell 75, 1203–1217.e5. 10.1016/j.molcel.2019.07.029.

54. Burke, J.M., Gilchrist, A.R., Sawyer, S.L., and Parker, R. (2021). RNase L limits host and viral protein synthesis via inhibition of mRNA export. Science Advances 7, eabh2479. 10.1126/sciadv.abh2479.

55. Donovan, J., Rath, S., Kolet-Mandrikov, D., and Korennykh, A. (2017). Rapid RNase L-driven arrest of protein synthesis in the dsRNA response without degradation of translation machinery. RNA 23, 1660–1671. 10.1261/rna.062000.117.

56. Whelan, J.N., Li, Y., Silverman, R.H., and Weiss, S.R. (2019). Zika Virus Production Is Resistant to RNase L Antiviral Activity. J Virol 93, e00313–19. 10.1128/JVI.00313-19.

57. Paul, D., and Bartenschlager, R. (2013). Architecture and biogenesis of plus-strand RNA virus replication factories. World J Virol 2, 32–48. 10.5501/wjv.v2.i2.32.

58. Silverman, R.H. (2007). Viral encounters with 2’,5’-oligoadenylate synthetase and RNase L during the interferon antiviral response. J Virol 81, 12720–12729. 10.1128/JVI.01471-07.

59. Schroeder, A., Mueller, O., Stocker, S., Salowsky, R., Leiber, M., Gassmann, M., Lightfoot, S., Menzel, W., Granzow, M., and Ragg, T. (2006). The RIN: an RNA integrity number for assigning integrity values to RNA measurements. BMC Molecular Biology 7, 3. 10.1186/1471-2199-7-3.

60. Karasik, A., Jones, G.D., DePass, A.V., and Guydosh, N.R. (2021). Activation of the antiviral factor RNase L triggers translation of non-coding mRNA sequences. Nucleic Acids Research 49, 6007–6026. 10.1093/nar/gkab036.

61. Schmidt, E.K., Clavarino, G., Ceppi, M., and Pierre, P. (2009). SUnSET, a nonradioactive method to monitor protein synthesis. Nat Methods 6, 275–277. 10.1038/nmeth.1314.

62. Ravi, V., Jain, A., Mishra, S., and Sundaresan, N.R. (2020). Measuring Protein Synthesis in Cultured Cells and Mouse Tissues Using the Non-radioactive SUnSET Assay. Current Protocols in Molecular Biology 133, e127. 10.1002/cpmb.127.

63. Santoro, M.G., Rossi, A., and Amici, C. (2003). NF-κB and virus infection: who controls whom. The EMBO Journal 22, 2552–2560. 10.1093/emboj/cdg267.

64. Mitchell, J.P., and Carmody, R.J. (2018). Chapter Two - NF-κB and the Transcriptional Control of Inflammation. In International Review of Cell and Molecular Biology Transcriptional Gene Regulation in Health and Disease., F. Loos, ed. (Academic Press), pp. 41–84. 10.1016/bs.ircmb.2017.07.007.

65. Lee, Y.-R., Kweon, S.-H., Kwon, K.-B., Park, J.-W., Yoon, T.-R., and Park, B.-H. (2009). Inhibition of IL-1β-mediated inflammatory responses by the IκBα super-repressor in human fibroblast-like synoviocytes. Biochemical and Biophysical Research Communications 378, 90–94. 10.1016/j.bbrc.2008.11.002.

66. Brown, K., Gerstberger, S., Carlson, L., Franzoso, G., and Siebenlist, U. (1995). Control of IκB-α Proteolysis by Site-Specific, Signal-Induced Phosphorylation. Science 267, 1485–1488. 10.1126/science.7878466.

67. Yan, X., Liu, Z., and Chen, Y. (2009). Regulation of TGF-β signaling by Smad7. Acta Biochimica et Biophysica Sinica 41, 263–272. 10.1093/abbs/gmp018.

68. Sobah, M.L., Liongue, C., and Ward, A.C. (2021). SOCS Proteins in Immunity, Inflammatory Diseases, and Immune-Related Cancer. Front Med (Lausanne) 8, 727987. 10.3389/fmed.2021.727987.

69. Dimitriou, I.D., Clemenza, L., Scotter, A.J., Chen, G., Guerra, F.M., and Rottapel, R. (2008). Putting out the fire: coordinated suppression of the innate and adaptive immune systems by SOCS1 and SOCS3 proteins. Immunological Reviews 224, 265–283. 10.1111/j.1600-065X.2008.00659.x.

70. Blumer, T., Coto-Llerena, M., Duong, F.H.T., and Heim, M.H. (2017). SOCS1 is an inducible negative regulator of interferon λ (IFN-λ)-induced gene expression in vivo. J Biol Chem 292, 17928–17938. 10.1074/jbc.M117.788877.

71. Bauer, S., Jendro, M.C., Wadle, A., Kleber, S., Stenner, F., Dinser, R., Reich, A., Faccin, E., Gödde, S., Dinges, H., et al. (2006). Fibroblast activation protein is expressed by rheumatoid myofibroblast-like synoviocytes. Arthritis Res Ther 8, R171. 10.1186/ar2080.

72. Zhang, H.E., Hamson, E.J., Koczorowska, M.M., Tholen, S., Chowdhury, S., Bailey, C.G., Lay, A.J., Twigg, S.M., Lee, Q., Roediger, B., et al. (2019). Identification of Novel Natural Substrates of Fibroblast Activation Protein-alpha by Differential Degradomics and Proteomics. Mol Cell Proteomics 18, 65–85. 10.1074/mcp.RA118.001046.

73. Fan, A., Wu, G., Wang, J., Lu, L., Wang, J., Wei, H., Sun, Y., Xu, Y., Mo, C., Zhang, X., et al. (2023). Inhibition of fibroblast activation protein ameliorates cartilage matrix degradation and osteoarthritis progression. Bone Res 11, 1–12. 10.1038/s41413-022-00243-8.

74. Croft, A.P., Campos, J., Jansen, K., Turner, J.D., Marshall, J., Attar, M., Savary, L., Wehmeyer, C., Naylor, A.J., Kemble, S., et al. (2019). Distinct fibroblast subsets drive inflammation and damage in arthritis. Nature 570, 246–251. 10.1038/s41586-019-1263-7.

75. Boyd, D.F., Allen, E.K., Randolph, A.G., Guo, X.J., Weng, Y., Sanders, C.J., Bajracharya, R., Lee, N.K., Guy, C.S., Vogel, P., et al. (2020). Exuberant fibroblast activity compromises lung function via ADAMTS-4. Nature 587, 466–471. 10.1038/s41586-020-2877-5.

76. Krausgruber, T., Fortelny, N., Fife-Gernedl, V., Senekowitsch, M., Schuster, L.C., Lercher, A., Nemc, A., Schmidl, C., Rendeiro, A.F., Bergthaler, A., et al. (2020). Structural cells are key regulators of organ-specific immune response. Nature 583, 296–302. 10.1038/s41586-020-2424-4.

77. Muta, D., Makino, K., Nakamura, H., Yano, S., Kudo, M., and Kuratsu, J.-I. (2011). Inhibition of eIF4E phosphorylation reduces cell growth and proliferation in primary central nervous system lymphoma cells. J Neurooncol 101, 33–39. 10.1007/s11060-010-0233-6.

78. Minnaert, A.-K., Vanluchene, H., Verbeke, R., Lentacker, I., De Smedt, S.C., Raemdonck, K., Sanders, N.N., and Remaut, K. (2021). Strategies for controlling the innate immune activity of conventional and self-amplifying mRNA therapeutics: Getting the message across. Adv Drug Deliv Rev 176, 113900. 10.1016/j.addr.2021.113900.

79. Frolova, E.I., Palchevska, O., Dominguez, F., and Frolov, I. Alphavirus-induced transcriptional and translational shutoffs play major roles in blocking the formation of stress granules. J Virol 97, e00979–23. 10.1128/jvi.00979-23.

80. Berglund, P., Finzi, D., Bennink, J.R., and Yewdell, J.W. (2007). Viral Alteration of Cellular Translational Machinery Increases Defective Ribosomal Products. Journal of Virology 81, 7220– 7229. 10.1128/jvi.00137-07.

81. Fros, J.J., and Pijlman, G.P. (2016). Alphavirus Infection: Host Cell Shut-Off and Inhibition of Antiviral Responses. Viruses 8, 166. 10.3390/v8060166.

82. Scheper, G.C., and Proud, C.G. (2002). Does phosphorylation of the cap-binding protein eIF4E play a role in translation initiation? European Journal of Biochemistry 269, 5350–5359. 10.1046/j.1432-1033.2002.03291.x.

83. Dyer, J.R., Michel, S., Lee, W., Castellucci, V.F., Wayne, N.L., and Sossin, W.S. (2003). An activity-dependent switch to cap-independent translation triggered by eIF4E dephosphorylation. Nat Neurosci 6, 219–220. 10.1038/nn1018.

84. Chang, C., Music, N., Cheung, M., Rossignol, E., Bedi, S., Patel, H., Safari, M., Lee, C., Otten, G.R., Settembre, E.C., et al. (2022). Self-amplifying mRNA bicistronic influenza vaccines raise cross-reactive immune responses in mice and prevent infection in ferrets. Molecular Therapy Methods & Clinical Development 27, 195–205. 10.1016/j.omtm.2022.09.013.

85. Gaucherand, L., and Gaglia, M.M. (2022). The Role of Viral RNA Degrading Factors in Shutoff of Host Gene Expression. Annu Rev Virol 9, 213–238. 10.1146/annurev-virology-100120-012345.

86. Xi, J., Snieckute, G., Martínez, J.F., Arendrup, F.S.W., Asthana, A., Gaughan, C., Lund, A.H., Bekker-Jensen, S., and Silverman, R.H. (2024). Initiation of a ZAKα-dependent ribotoxic stress response by the innate immunity endoribonuclease RNase L. Cell Reports 43. 10.1016/j.celrep.2024.113998.

87. Guydosh, N.R., Kimmig, P., Walter, P., and Green, R. (2017). Regulated Ire1-dependent mRNA decay requires no-go mRNA degradation to maintain endoplasmic reticulum homeostasis in S. pombe. eLife 6, e29216. 10.7554/eLife.29216.

88. Snieckute, G., Genzor, A.V., Vind, A.C., Ryder, L., Stoneley, M., Chamois, S., Dreos, R., Nordgaard, C., Sass, F., Blasius, M., et al. (2022). Ribosome stalling is a signal for metabolic regulation by the ribotoxic stress response. Cell Metab 34, 2036–2046.e8. 10.1016/j.cmet.2022.10.011.

89. Li, G., Xiang, Y., Sabapathy, K., and Silverman, R.H. (2004). An Apoptotic Signaling Pathway in the Interferon Antiviral Response Mediated by RNase L and c-Jun NH2-terminal Kinase *. Journal of Biological Chemistry 279, 1123–1131. 10.1074/jbc.M305893200.

90. Roretz, C. von, and Gallouzi, I.-E. (2010). Protein Kinase RNA/FADD/Caspase-8 Pathway Mediates the Proapoptotic Activity of the RNA-binding Protein Human Antigen R (HuR). The Journal of Biological Chemistry 285, 16806. 10.1074/jbc.M109.087320.

91. Zuo, W., Wakimoto, M., Kozaiwa, N., Shirasaka, Y., Oh, S.-W., Fujiwara, S., Miyachi, H., Kogure, A., Kato, H., and Fujita, T. (2022). PKR and TLR3 trigger distinct signals that coordinate the induction of antiviral apoptosis. Cell Death Dis 13, 1–15. 10.1038/s41419-022-05101-3.

92. Xu, C., Gamil, A.A.A., Munang’andu, H.M., and Evensen, Ø. (2018). Apoptosis Induction by dsRNA-Dependent Protein Kinase R (PKR) in EPC Cells via Caspase 8 and 9 Pathways. Viruses 10, 526. 10.3390/v10100526.

93. Gil, J., and Esteban, M. (2000). The interferon-induced protein kinase (PKR), triggers apoptosis through FADD-mediated activation of caspase 8 in a manner independent of Fas and TNF-α receptors. Oncogene 19, 3665–3674. 10.1038/sj.onc.1203710.

94. Zhang, P., and Samuel, C.E. (2007). Protein Kinase PKR Plays a Stimulus-and Virus-Dependent Role in Apoptotic Death and Virus Multiplication in Human Cells. Journal of Virology 81, 8192–8200. 10.1128/jvi.00426-07.

95. Leitner, W.W., Hwang, L.N., Bergmann-Leitner, E.S., Finkelstein, S.E., Frank, S., and Restifo, N.P. (2004). Apoptosis is essential for the increased efficacy of alphaviral replicase-based DNA vaccines. Vaccine 22, 1537–1544. 10.1016/j.vaccine.2003.10.013.

96. Wu, Y.-Y., Sun, T.-K., Chen, M.-S., Munir, M., and Liu, H.-J. (2023). Oncolytic viruses-modulated immunogenic cell death, apoptosis and autophagy linking to virotherapy and cancer immune response. Front. Cell. Infect. Microbiol. 13. 10.3389/fcimb.2023.1142172.

97. Iordanov, M.S., Wong, J., Bell, J.C., and Magun, B.E. (2001). Activation of NF-κB by Double-Stranded RNA (dsRNA) in the Absence of Protein Kinase R and RNase L Demonstrates the Existence of Two Separate dsRNA-Triggered Antiviral Programs. Molecular and Cellular Biology 21, 61–72. 10.1128/MCB.21.1.61-72.2001.

98. Herdy, B., Jaramillo, M., Svitkin, Y.V., Rosenfeld, A.B., Kobayashi, M., Walsh, D., Alain, T., Sean, P., Robichaud, N., Topisirovic, I., et al. (2012). Translational control of the activation of transcription factor NF-κB and production of type I interferon by phosphorylation of the translation factor eIF4E. Nat Immunol 13, 543–550. 10.1038/ni.2291.

99. Liu, T., Zhang, L., Joo, D., and Sun, S.-C. (2017). NF-κB signaling in inflammation. Sig Transduct Target Ther 2, 1–9. 10.1038/sigtrans.2017.23.

100. Luo, J.-L., Kamata, H., and Karin, M. (2005). IKK/NF-κB signaling: balancing life and death – a new approach to cancer therapy. Journal of Clinical Investigation 115, 2625–2632. 10.1172/JCI26322.

101. Miagkov, A.V., Kovalenko, D.V., Brown, C.E., Didsbury, J.R., Cogswell, J.P., Stimpson, S.A., Baldwin, A.S., and Makarov, S.S. (1998). NF-κB activation provides the potential link between inflammation and hyperplasia in the arthritic joint. Proceedings of the National Academy of Sciences 95, 13859–13864. 10.1073/pnas.95.23.13859.

102. Zhang, H.G., Huang, N., Liu, D., Bilbao, L., Zhang, X., Yang, P., Zhou, T., Curiel, D.T., and Mountz, J.D. (2000). Gene therapy that inhibits nuclear translocation of nuclear factor kappaB results in tumor necrosis factor alpha-induced apoptosis of human synovial fibroblasts. Arthritis Rheum 43, 1094–1105. 10.1002/1529-0131(200005)43:5<1094::AID-ANR20>3.0.CO;2-V.

103. Xu, Y., Bialik, S., Jones, B.E., Iimuro, Y., Kitsis, R.N., Srinivasan, A., Brenner, D.A., and Czaja, M.J. (1998). NF-kappaB inactivation converts a hepatocyte cell line TNF-alpha response from proliferation to apoptosis. Am J Physiol 275, C1058–1066. 10.1152/ajpcell.1998.275.4.C1058.

104. Sago, C.D., Krupczak, B.R., Lokugamage, M.P., Gan, Z., and Dahlman, J.E. (2019). Cell Subtypes Within the Liver Microenvironment Differentially Interact with Lipid Nanoparticles. Cellular and Molecular Bioengineering 12, 389. 10.1007/s12195-019-00573-4.

105. Yamamura, Y., Hua, X., Bergelson, S., and Lodish, H.F. (2000). Critical Role of Smads and AP-1 Complex in Transforming Growth Factor-β-dependent Apoptosis. Journal of Biological Chemistry 275, 36295–36302. 10.1074/jbc.M006023200.

106. Patil, S., Wildey, G.M., Brown, T.L., Choy, L., Derynck, R., and Howe, P.H. (2000). Smad7 is induced by CD40 and protects WEHI 231 B-lymphocytes from transforming growth factor-beta - induced growth inhibition and apoptosis. J Biol Chem 275, 38363–38370. 10.1074/jbc.M004861200.

107. Zitzmann, K., Brand, S., De Toni, E.N., Baehs, S., Göke, B., Meinecke, J., Spöttl, G., Meyer, H.H.H.D., and Auernhammer, C.J. (2007). SOCS1 Silencing Enhances Antitumor Activity of Type I IFNs by Regulating Apoptosis in Neuroendocrine Tumor Cells. Cancer Research 67, 5025–5032. 10.1158/0008-5472.CAN-06-2575.

108. Mann-Chandler, M.N., Kashyap, M., Wright, H.V., Norozian, F., Barnstein, B.O., Gingras, S., Parganas, E., and Ryan, J.J. (2005). IFN-γ Induces Apoptosis in Developing Mast Cells1. The Journal of Immunology 175, 3000–3005. 10.4049/jimmunol.175.5.3000.

109. Kimura, A., Naka, T., Nagata, S., Kawase, I., and Kishimoto, T. (2004). SOCS-1 suppresses TNF-alpha-induced apoptosis through the regulation of Jak activation. Int Immunol 16, 991–999. 10.1093/intimm/dxh102.

110. Madonna, S., Scarponi, C., Pallotta, S., Cavani, A., and Albanesi, C. (2012). Anti-apoptotic effects of suppressor of cytokine signaling 3 and 1 in psoriasis. Cell Death Dis 3, e334–e334. 10.1038/cddis.2012.69.

111. Yan, L., Tang, Q., Shen, D., Peng, S., Zheng, Q., Guo, H., Jiang, M., and Deng, W. (2008). SOCS-1 Inhibits TNF-α-Induced Cardiomyocyte Apoptosis via ERK1/2 Pathway Activation. Inflammation 31, 180–188. 10.1007/s10753-008-9063-5.

112. Goker Bagca, B., and Biray Avci, C. (2020). The potential of JAK/STAT pathway inhibition by ruxolitinib in the treatment of COVID-19. Cytokine & Growth Factor Reviews 54, 51–61. 10.1016/j.cytogfr.2020.06.013.

113. Liau, N.P.D., Laktyushin, A., Lucet, I.S., Murphy, J.M., Yao, S., Whitlock, E., Callaghan, K., Nicola, N.A., Kershaw, N.J., and Babon, J.J. (2018). The molecular basis of JAK/STAT inhibition by SOCS1. Nat Commun 9, 1558. 10.1038/s41467-018-04013-1.

114. Skidmore, A.M., Adcock, R.S., Jonsson, C.B., Golden, J.E., and Chung, D.-H. (2020). Benzamidine ML336 inhibits plus and minus strand RNA synthesis of Venezuelan equine encephalitis virus without affecting host RNA production. Antiviral Research 174, 104674. 10.1016/j.antiviral.2019.104674.

115. Schroeder, C.E., Yao, T., Sotsky, J., Smith, R.A., Roy, S., Chu, Y.-K., Guo, H., Tower, N.A., Noah, J.W., McKellip, S., et al. (2014). Development of (E)-2-((1,4-Dimethylpiperazin-2-ylidene)amino)-5-nitro-N-phenylbenzamide, ML336: Novel 2-Amidinophenylbenzamides as Potent Inhibitors of Venezuelan Equine Encephalitis Virus. J. Med. Chem. 57, 8608–8621. 10.1021/jm501203v.

116. Tregoning, J.S., Stirling, D.C., Wang, Z., Flight, K.E., Brown, J.C., Blakney, A.K., McKay, P.F., Cunliffe, R.F., Murugaiah, V., Fox, C.B., et al. (2023). Formulation, inflammation, and RNA sensing impact the immunogenicity of self-amplifying RNA vaccines. Molecular Therapy Nucleic Acids 31, 29–42. 10.1016/j.omtn.2022.11.024.

117. Bathula, N.V., Friesen, J.J., Casmil, I.C., Wayne, C.J., Liao, S., Soriano, S.K.V., Ho, C.H., Strumpel, A., and Blakney, A.K. (2024). Delivery vehicle and route of administration influences self-amplifying RNA biodistribution, expression kinetics, and reactogenicity. Journal of Controlled Release 374, 28–38. 10.1016/j.jconrel.2024.07.078.

118. Oda, Y., Kumagai, Y., Kanai, M., Iwama, Y., Okura, I., Minamida, T., Yagi, Y., Kurosawa, T., Greener, B., Zhang, Y., et al. (2024). Immunogenicity and safety of a booster dose of a self-amplifying RNA COVID-19 vaccine (ARCT-154) versus BNT162b2 mRNA COVID-19 vaccine: a double-blind, multicentre, randomised, controlled, phase 3, non-inferiority trial. The Lancet Infectious Diseases 24, 351–360. 10.1016/S1473-3099(23)00650-3.

119. Saraf, A., Gurjar, R., Kaviraj, S., Kulkarni, A., Kumar, D., Kulkarni, R., Virkar, R., Krishnan, J., Yadav, A., Baranwal, E., et al. (2024). An Omicron-specific, self-amplifying mRNA booster vaccine for COVID-19: a phase 2/3 randomized trial. Nat Med 30, 1363–1372. 10.1038/s41591-024-02955-2.

120. Roman-Blas, J.A., and Jimenez, S.A. (2006). NF-κB as a potential therapeutic target in osteoarthritis and rheumatoid arthritis. Osteoarthritis and Cartilage 14, 839–848. 10.1016/j.joca.2006.04.008.

121. Shen, J., Li, S., and Chen, D. (2014). TGF-β signaling and the development of osteoarthritis. Bone Res 2, 1–7. 10.1038/boneres.2014.2.

122. Zhou, Q., Ren, Q., Jiao, L., Huang, J., Yi, J., Chen, J., Lai, J., Ji, G., and Zheng, T. (2022). The potential roles of JAK/STAT signaling in the progression of osteoarthritis. Front. Endocrinol. 13. 10.3389/fendo.2022.1069057.

123. Blackwell, N.M., Sembi, P., Newson, J.S., Lawrence, T., Gilroy, D.W., and Kabouridis, P.S. (2004). Reduced infiltration and increased apoptosis of leukocytes at sites of inflammation by systemic administration of a membrane-permeable IκBα repressor. Arthritis & Rheumatism 50, 2675–2684. 10.1002/art.20467.

124. Chen, S.-Y., Shiau, A.-L., Wu, C.-L., and Wang, C.-R. (2016). Intraarticular overexpression of Smad7 ameliorates experimental arthritis. Sci Rep 6, 35163. 10.1038/srep35163.

125. Ha, Y.-J., Choi, Y.S., Kang, E.H., Shin, K., Kim, T.K., Song, Y.W., and Lee, Y.J. (2016). SOCS1 suppresses IL-1β-induced C/EBPβ expression via transcriptional regulation in human chondrocytes. Exp Mol Med 48, e241–e241. 10.1038/emm.2016.47.

126. Freppel, W., Lim, E.X.Y., Rudd, P.A., and Herrero, L.J. (2024). Synoviocytes assist in modulating the effect of Ross River virus infection in micromass-cultured primary human chondrocytes. J Med Microbiol 73, 001859. 10.1099/jmm.0.001859.

127. Legros, V., Belarbi, E., Jeannin, P., Kümmerer, B., Hardy, D., Desprès, P., Gessain, A., Roques, P., Ceccaldi, P.-E., and Choumet, V. (2024). A recombinant CHIKV-NLuc virus identifies chondrocytes as target of Chikungunya virus in a immunocompetent mouse model. Preprint at bioRxiv, 10.1101/2024.05.20.594924.

128. Wäldele, S., Koers-Wunrau, C., Beckmann, D., Korb-Pap, A., Wehmeyer, C., Pap, T., and Dankbar, B. (2015). Deficiency of fibroblast activation protein alpha ameliorates cartilage destruction in inflammatory destructive arthritis. Arthritis Res Ther 17, 12. 10.1186/s13075-015-0524-6.

129. Hồ, N.T., Hughes, S.G., Ta, V.T., Phan, L.T., Đỗ, Q., Nguyễn, T.V., Phạm, A.T.V., Thị Ngọc Đặng, M., Nguyễn, L.V., Trịnh, Q.V., et al. (2024). Safety, immunogenicity and efficacy of the self-amplifying mRNA ARCT-154 COVID-19 vaccine: pooled phase 1, 2, 3a and 3b randomized, controlled trials. Nat Commun 15, 4081. 10.1038/s41467-024-47905-1.

130. Kimura, T., Leal, J.M., Simpson, A., Warner, N.L., Berube, B.J., Archer, J.F., Park, S., Kurtz, R., Hinkley, T., Nicholes, K., et al. (2023). A localizing nanocarrier formulation enables multi-target immune responses to multivalent replicating RNA with limited systemic inflammation. Molecular Therapy 31, 2360–2375. 10.1016/j.ymthe.2023.06.017.

131. Gadella, T.W.J., van Weeren, L., Stouthamer, J., Hink, M.A., Wolters, A.H.G., Giepmans, B.N.G., Aumonier, S., Dupuy, J., and Royant, A. (2023). mScarlet3: a brilliant and fast-maturing red fluorescent protein. Nat Methods 20, 541–545. 10.1038/s41592-023-01809-y.

132. Costantini, L.M., Baloban, M., Markwardt, M.L., Rizzo, M.A., Guo, F., Verkhusha, V.V., and Snapp, E.L. (2015). A palette of fluorescent proteins optimized for diverse cellular environments. Nat Commun 6, 7670. 10.1038/ncomms8670.

133. Floor, S. (2019). Denaturing agarose gels for large RNAs with glyoxal.

134. Rio, D.C. (2015). Denaturation and Electrophoresis of RNA with Glyoxal. Cold Spring Harb Protoc 2015, pdb.prot081000. 10.1101/pdb.prot081000.

135. Chakrabarti, S., Hore, Z., Pattison, L.A., Lalnunhlimi, S., Bhebhe, C.N., Callejo, G., Bulmer, D.C., Taams, L.S., Denk, F., and Smith, E.S.J. (2020). Sensitization of knee-innervating sensory neurons by tumor necrosis factor-α-activated fibroblast-like synoviocytes: an in vitro, coculture model of inflammatory pain. Pain 161, 2129–2141. 10.1097/j.pain.0000000000001890.

136. Li, C.H., and Lee, C.K. (1993). Minimum cross entropy thresholding. Pattern Recognition 26, 617– 625. 10.1016/0031-3203(93)90115-D.

